# Comprehensive identification of somatic nucleotide variants in human brain tissue

**DOI:** 10.1101/2020.10.10.332213

**Authors:** Yifan Wang, Taejeong Bae, Jeremy Thorpe, Maxwell A. Sherman, Attila G. Jones, Sean Cho, Kenneth Daily, Yanmei Dou, Javier Ganz, Alon Galor, Irene Lobon, Reenal Pattni, Chaggai Rosenbluh, Simone Tomasi, Livia Tomasini, Xiaoxu Yang, Bo Zhou, Schahram Akbarian, Laurel L. Ball, Sara Bizzotto, Sarah B. Emery, Ryan Doan, Liana Fasching, Yeongjun Jang, David Juan, Esther Lizano, John B. Moldovan, Rujuta Narurkar, Matthew T. Oetjens, Shobana Sekar, Joo Heon Shin, Eduardo Soriano, Richard E. Straub, Weichen Zhou, Andrew Chess, Joseph G. Gleeson, Tomas Marquès-Bonet, Peter J. Park, Mette A. Peters, Jonathan Pevsner, Christopher A. Walsh, Daniel R. Weinberger, Brain Somatic Mosaicism Network, Flora M. Vaccarino, John V. Moran, Alexander E. Urban, Jeffrey M. Kidd, Ryan E. Mills, Alexej Abyzov

## Abstract

Post-zygotic mutations incurred during DNA replication, DNA repair, and other cellular processes lead to somatic mosaicism. Somatic mosaicism is an established cause of various diseases, including cancers. However, detecting mosaic variants in DNA from non-cancerous somatic tissues poses significant challenges, particularly if the variants only are present in a small fraction of cells. Here, the Brain Somatic Mosaicism Network conducted a coordinated, multi-institutional study to: (i) examine the ability of existing methods to detect simulated somatic single nucleotide variants (SNVs) in DNA mixing experiments; (ii) generate multiple replicates of whole genome sequencing data from the dorsolateral prefrontal cortex, other brain regions, dura mater, and dural fibroblasts of a single neurotypical individual; (iii) devise strategies to discover somatic SNVs; and (iv) apply various approaches to validate somatic SNVs. These efforts led to the identification of 43 *bona fide* somatic SNVs that ranged in variant allele fractions from ~0.005 to ~0.28. Guided by these results, we devised best practices for calling mosaic SNVs from 250X whole genome sequencing data in the accessible portion of the human genome that achieve 90% specificity and sensitivity. Finally, we demonstrated that analysis of multiple bulk DNA samples from a single individual allows the reconstruction of early developmental cell lineage trees. Thus, this study provides a unified set of best practices to detect somatic SNVs in non-cancerous tissues. The data and methods are freely available to the scientific community and should serve as a guide to assess the contributions of somatic SNVs to neuropsychiatric diseases.

## Introduction

Genomic sequence variants may be inherited vertically (*i.e.*, transmitted through the germline) or generated after zygote formation (*i.e.*, leading to somatic or gonadal mosaicism). It is well established that somatic mosaicism occurs in cells of phenotypically normal individuals^1,2,3,4,5,6,7,8,9,10,11,12,13,14,15,16,17^ and can lead to various diseases^18^. However, the prevalence of somatic mosaicism and the extent to which it contributes to diseases outside of cancers requires elucidation^18^.

Recent studies have estimated that each cell within the human brain contains hundreds to a few thousand somatic single nucleotide variants (SNVs) and that a smaller fraction of cells harbor somatic copy number variations (CNVs) and mobile genetic element (*i.e.*, retrotransposon) insertions^10,15,17,19,20,21,22^. Dozens of somatic SNVs are present at high variant allele fractions (VAFs) across multiple tissues, indicating that they arose during early development^17,23^. By comparison, some somatic SNVs are present at low VAFs in a small number of cells and have limited tissue distributions, suggesting they arose later in development^15,16,17^.

Single cell DNA sequencing is the most direct approach to identify somatic variants. However, mutations introduced during the amplification and/or generation of single cell DNA sequencing libraries, as well as nonuniform DNA amplification biases, make it difficult to discriminate *bona fide* mosaic SNVs from procedural artifacts^24^. Moreover, this approach for identifying mosaic SNVs requires sampling a large number of cells in a given individual and, consequently, is cost intensive.

Another approach to identify mosaic variants involves comparing bulk cell populations from two tissue samples derived from the same individual — the sample of interest and a control sample — as is performed routinely during the analysis of cancer genomes. However, this approach is limited by the inability to define a proper control tissue because mosaic SNVs, particularly ones that arise during early development, are often present in multiple tissues across the body. Thus, the development of a unified set of best practices to detect somatic SNVs from bulk whole genome sequencing (WGS) datasets would provide an alternative, cost-effective approach to identify somatic SNVs.

In this study, members of the Brain Somatic Mosaicism Network (BSMN) conducted a coordinated, multi-institutional study that analyzed mosaicism in a single neurotypical brain sample and established unified standards for calling and validating mosaic SNVs from bulk WGS and WES data.

## Results

### The detection of simulated somatic SNVs in DNA mixing experiments

We first assessed the ability of three variant callers commonly used to detect germline single nucleotide polymorphisms (SNPs) and somatic SNVs in cancers (*i.e.*, the GATK HaplotypeCaller^25^, MuTect2^26^, and Strelka2^27^) to detect simulated somatic SNVs. We mixed genomic DNAs derived from transformed lymphoblastoid cell lines of four unrelated individuals at different proportions (Figs. 1a **& S1**; see Mix 1 and Mix 2), sequenced the resultant mixtures to ~100X coverage, and assessed the ability of the callers to detect the simulated somatic SNVs (Figs. 1b **& S2**). In general, none of the callers were sensitive enough to reliably detect SNVs present at low (<0.10) VAFs. Combining the Mix 1 and Mix 2 datasets to double sequencing coverage only marginally improved sensitivity (Figs. 1b **& S2**). Moreover, MuTect2 and Strelka2, which are designed to detect somatic SNVs that are present in tumors but not matched normal samples, lacked the sensitivity to detect simulated mosaic SNVs shared between two matched samples (Figs. 1b **& S2**).

**Figure 1:**
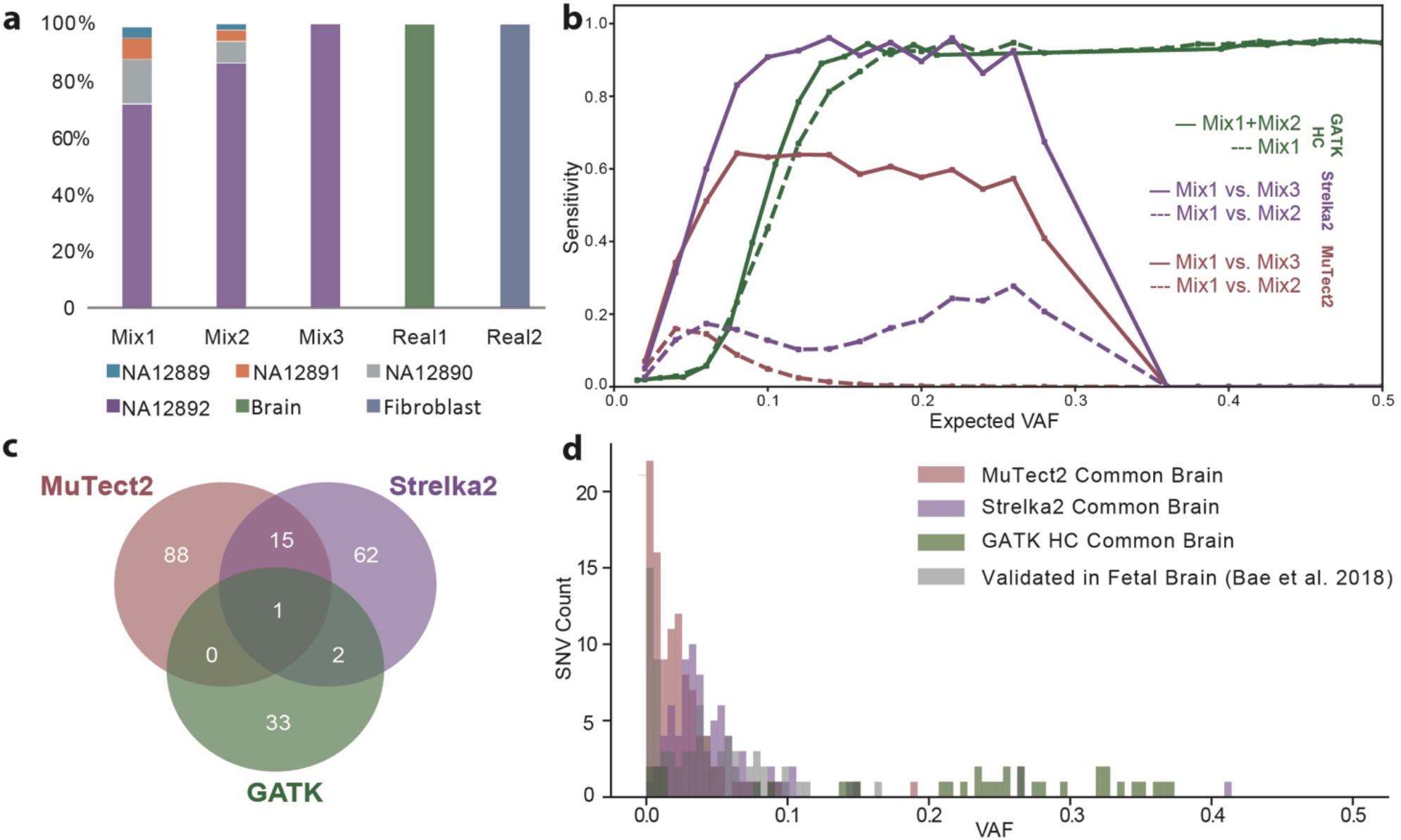
Assessment of existing tools to detect simulated mosaic SNVs in DNA mixing experiments (a & b) or candidate somatic SNVs in the common reference brain sample (c & d). (*a*) Genomic DNAs from four commonly-used human lymphoblastoid cell lines were mixed at various proportions (x-axis) and subjected to WGS; germline SNPs from the cell lines are present at a range of allele frequencies (y-axis) in the different mixes and act as a proxy for mosaic SNVs. (*b*) VAF of simulated SNVs (x-axis) *vs*. sensitivity of detection (y-axis) for the three described SNV callers. (*c*) A Venn diagram demonstrating that existing tools are widely discordant in their ability to call mosaic SNVs present in the common brain sample. (*d*) The distribution of candidate SNV VAFs (x-axis) and the numbers of candidate mosaic SNV calls (y-axis) detected by existing tools.

We next compared the performance of the three variant callers to detect putative somatic SNVs in ~250X coverage WGS data derived from post-mortem dorsolateral prefrontal cortex (DLPFC; herein called the common reference brain; see below) and dural fibroblasts derived from a single neurotypical individual. We discarded likely germline variants having ~0.5 VAFs and observed that the remaining putative somatic SNVs exhibited little overlap among the three callers (Fig. 1c). The VAF distributions identified using the variant callers also differed significantly from validated mosaic variants previously reported in fetal brain^17^ (Fig. 1d). Thus, the naive application of existing callers appear to lack the sensitivity and precision necessary to robustly call somatic SNVs from bulk brain tissue, indicating a need for new approaches to address this challenge.

### The generation of a somatic SNV call set from a neurotypical brain sample

To arrive at an unbiased approach to confidently identify somatic SNVs, we dissected a DLPFC sample from the reference brain of a neurotypical individual, pulverized it to enhance homogeneity, and distributed aliquots of this sample, as well as aliquots of a matched isogenic primary dural fibroblast cell line, to individual BSMN working groups. Each BSMN group independently performed DNA extractions and generated either whole genome (four datasets ranging from ~85X to ~245X sequencing coverage) or exome sequencing (two datasets of ~350X and ~435X sequencing coverage) data (Fig. 2a; Replicates 1 through 6). The resultant reads were uniformly processed using a common workflow (see **Methods**) and mapped to the human genome (version GRCh37d5) (**Fig. S3**). Group-specific computational approaches and filtering strategies then were used to identify putative somatic SNVs (Fig. 2a; Methods 1 through 6).

**Figure 2:**
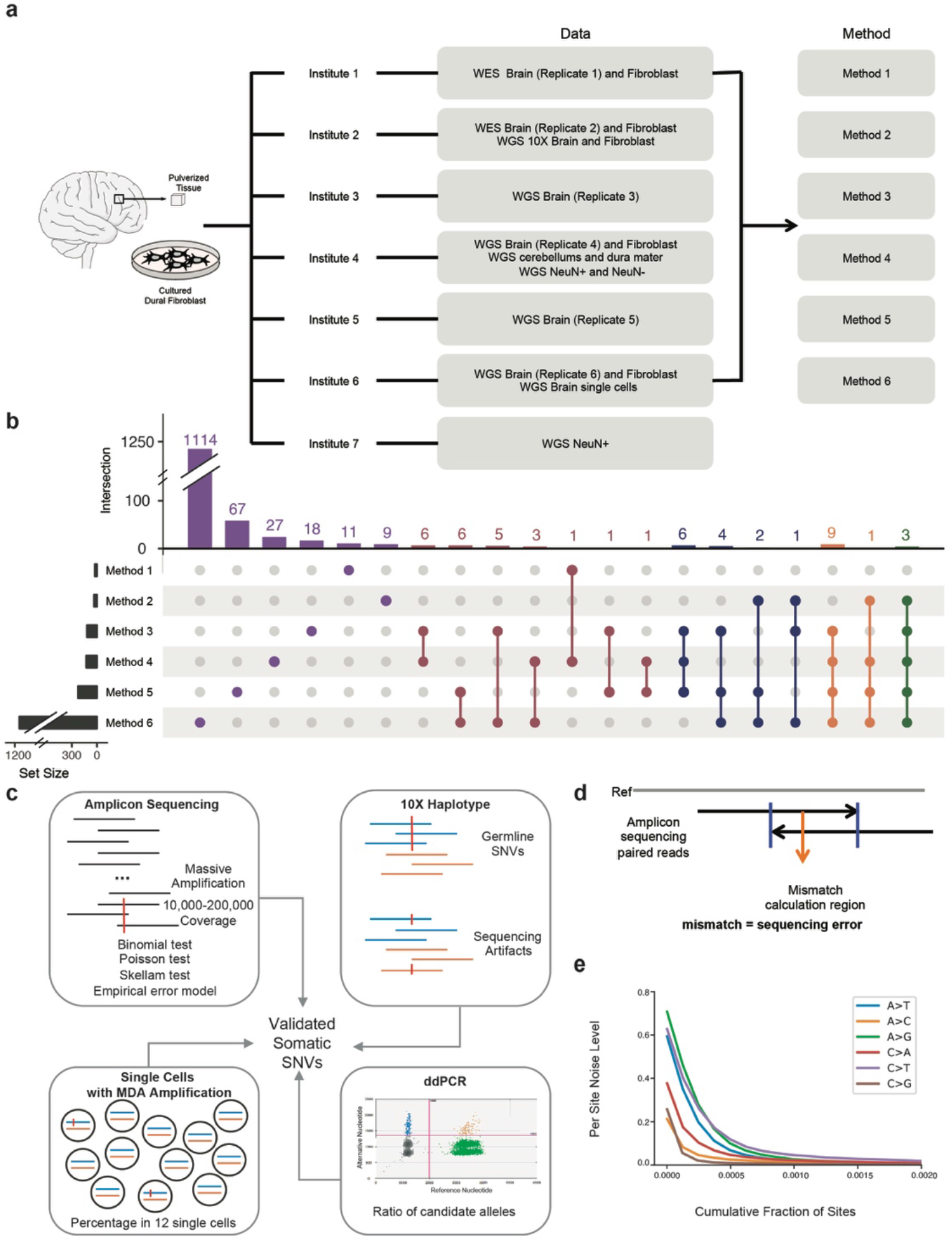
Overview of the mosaic SNV discovery and validation pipeline. (*a*) WGS or WES datasets were generated by six BSMN working groups using a commonly-shared, homogenized DLPFC sample from a neurotypical individual and isogenic dural fibroblasts. Six different analytical methods initially were used to call mosaic SNVs. WGS data also was generated from sorted NeuN+ and NeuN-cells from DLPFC, cerebellum, and dura mater samples. Chromium 10X linked-read sequencing data was generated from DLPFC and dural fibroblast samples. Single-cell WGS sequencing was conducted on twelve NeuN+ neurons from the DLPFC. These datasets were used to validate mosaic SNVs. (*b*) Overlap of putative mosaic SNV calls using different analytical approaches. Indicated are the numbers of mosaic SNV calls (x-axis) and the numbers of mosaic SNV calls identified using different analytical approaches (y-axis; circles with connecting lines indicate candidate SNVs identified by multiple approaches). (*c*) Candidate SNVs were subject to validation experiments using four complementary approaches. (*d*) Rationale of the empirical substitution error model applied to validate mosaic SNVs in PCR amplicon-based sequencing experiments. (*e*) An example of the empirical nucleotide error profiles encountered in a PCR amplicon-based sequencing experiment. Shown is the cumulative fraction of sites (x-axis) and per site noise levels (y-axis).

The most widely used WGS technologies result in relatively short (~100-150 bps) sequence reads that are derived from the ends of small (typically of 350-450 bps) DNA fragments. As such, many reads do not unambiguously map to a single genomic locus. For example, it is difficult to confidently assign short DNA reads to evolutionarily young and/or human-specific retrotransposon-derived sequences (*e.g.*, LINE-1 and Alu elements)^28,29^. Similarly, segmental duplications sharing high sequence identity, as well as tandem repeat sequences present in centromeric, telomeric, and subtelomeric genomic regions pose significant mapping challenges^30^. Thus, we applied the 1000 Genomes Project Strict Mask as a filter, which identifies positions in the human reference where sequence coverage does not significantly deviate from the expected average values across human populations. This mask defined ~73% of the human genome as accessible and each BSMN working group only called putative mosaic SNVs in the accessible fraction of the human genome.

The BSMN working groups used different analytical strategies to identify putative somatic SNVs (*e.g.*, single sample calling, paired sample calling, or a combination of both strategies, as well as various filtering strategies; see **Methods**). Some groups opted for higher precision calling approaches, yielding fewer, but presumably more precise mosaic SNV call sets. Other groups casted a wider net and opted to generate more inclusive, but presumably less precise call sets. Consequently, the initial numbers of putative somatic SNV calls were quite discordant and varied significantly among the working groups. For example, none of the putative SNV calls were identified by each of the six working groups and only three calls were identified by five of six groups (Figs. 2b **Table S1**).

We next assessed the sensitivity and specificity of the filters used by different BSMN working groups to generate individual call sets. First, we prioritized the 1114 putative mosaic SNV calls into four categories using the strategy outlined in Table 1. We reasoned that putative mosaic SNVs called by multiple groups using different approaches and from multiple data sources/replicates (Multi-call SNVs) would have the highest probability of yielding *bona fide* somatic SNVs. By comparison, SNVs identified using one approach from different data sources (Approach Singletons) and SNVs identified using multiple approaches from one data source (Data Source Singletons) might be expected to have more false-positive calls when compared to Multi-call SNVs. Finally, SNVs identified using a single approach from a single data source (Absolute Singletons) might be expected to have the fewest number of true mosaic SNV calls.

**Table 1.** Categories of Validated Mosaic SNVs

| Categories | Definition | Number of Mosaic SNVs selected for validation (out of total) | Number of Validated Mosaic SNVs |
| --- | --- | --- | --- |
| Multi-calls | Identified by multiple approaches, supportive evidence from multiple data sources | 45 out of 45 | 33 |
| Approach Singletons | Identified by one approach, supportive evidence from multiple data sources | 311 out of 1101 | 10 |
| Data Source Singletons | Identified by multiple approaches, supportive evidence from one data source | 4 out of 4 | 0 |
| Absolute Singletons | Identified by one approach, supportive evidence from one data source | 40 out of 148 | 0 |

We subjected 400 putative mosaic SNVs to validation experiments (**Tables S1 & S2**). The validation set included all the putative SNVs detected using two or more approaches (Data Source Singletons and Multi-calls; 49 calls), all of the calls detected individually using Methods 1-5 (Approach and Absolute Singletons; 124 calls), and a subset of randomly chosen calls detected individually by Method 6, which had the largest number of calls (Approach and Absolute Singletons; 227 calls). We divided the putative mosaic SNV calls into five sets of 100 variants; 20% of the SNV calls overlapped between the sets. The putative calls sets then were distributed to four BSMN working groups for PCR amplicon-based deep sequencing validation (Fig. 2c). Another group attempted to validate all 400 sites using a multiplex PCR-based targeted single-end re-sequencing assay (see **Methods**). Together, these strategies allowed testing of 272 putative SNVs by two independent working groups.

To aid in the validation of putative somatic SNVs we conducted the following analyses: (i) whenever possible, we used the 3, overlapping paired-end reads derived from each PCR amplicon to correct for sequencing errors (Fig. 2d); and (ii) we derived a nucleotide substitution error profile for each PCR amplicon-based sequencing library (Fig. 2e). To identify and eliminate systematic DNA amplification and sequencing artifacts, 4/5 groups also sequenced the same PCR validation amplicons from genomic DNA derived from the control NA12878 lymphoblastoid cell line^31^. This process yielded 233 concordant and 39 discordant putative validation calls. The majority (22/39) of discordant calls represented mosaic SNVs present at low VAFs (<0.01).

We resolved the discordant mosaic SNV calls by prioritizing conclusions from amplicon-seq experiments with overlapping paired reads, which provided higher confidence base calls within mapping reads. Additionally, we considered orthogonal experimental evidence (*i.e.*. whether the putative mosaic SNV was detected as an alternative read on a single haplotype in a ~70X coverage Chromium 10X linked-read dataset and/or whether the putative mosaic SNV was detected in NeuN+ single cell DNA sequencing datasets generated from the common reference sample) (Fig. 2c; see **Methods**). Finally, to verify the validation status, we performed droplet digital PCR (ddPCR) experiments for thirteen calls (**Table S3**). These calls included a germline SNP within a heterozygous genomic duplication, three false-positive calls, and nine previously validated mosaic SNVs. The ddPCR experiments demonstrated remarkable consistency with the VAFs determined using the PCR amplicon-based sequencing experiments, confirmed the predicted ~0.3 VAF for the germline SNP within a genomic duplication, invalidated the false-positive calls, and led to the re-validation of eight mosaic SNVs. One mosaic SNV validated by amplicon sequencing with a low VAF (0.0053) was not validated using ddPCR, highlighting potential limitations of PCR amplicon-based sequencing and/or ddPCR to validate mosaic SNVs at very low VAFs.

In total, 43 mosaic SNVs were validated as *bona fide* somatic SNV calls. The validated mosaic SNVs were present at VAFs that ranged from ~0.005 to ~0.28. None of the mosaic SNVs occurred within coding exons, one (chr19:9,493,288 G>A) was present in a non-coding exon (*i.e.*, 3′-UTR of *ZNF177)*, and one intronic variant (chr8:103,281,483 C>T) also had supportive reads in WES datasets, which only effectively survey 1%-2% of the human genome. Moreover, consistent with previous data^17^, the SNV mutation spectrum was dominated by C>T transitions (**Fig. S4**). The filtering strategies, sources of false-positive calls, and caveats when conducting variant discovery and validation experiments are discussed in greater detail below.

### Sources of false-positive SNV calls and filtering strategies

We sought to understand properties that distinguish *bona fide* somatic SNV calls from false-positive calls. Additional criteria used to identify and eliminate false-positive calls included: (i) assessing the base quality score of reads supporting the candidate SNV; (ii) identifying imbalances in candidate SNV counts in forward *vs.* reverse sequencing reads; (iii) examining local assemblies of candidate somatic SNVs present within or near polymorphic insertion/deletion mutations (indels), homopolymeric tracts, and structural variants; and (iv) determining whether the candidate SNV arose on a single parental haplotype. Below we briefly summarize the sources of false-positive calls discovered in our study.

### Germline SNPs

The PCR amplicon-based validation experiments coupled with the availability of Chromium 10X linked-read data, single cell DNA sequencing datasets, and germline copy number assessments in the common reference brain sample helped identify putative mosaic SNVs present at VAFs of <0.50 that ultimately were determined to be germline SNPs. For example, our initial analyses identified 29 putative SNVs that appeared at <0.50 VAFs; however, PCR amplicon-based sequencing validation demonstrated that these variants were present at ~0.50 VAFs. Consistently, Chromium 10X linked-read data demonstrated that 18/29 variants were present on a single parental haplotype (“2 haplotype” calls) and 25/29 variants were present in at least six of twelve single cell sequencing datasets (Fig. 3). Finally, 14/29 putative mosaic SNVs were located within regions of the genome containing germline copy number gains, leading to <0.50 VAFs (**Fig. S5**). This latter result highlights the importance of performing copy number characterization on experimental samples prior to calling mosaic SNVs.

**Figure 3:**
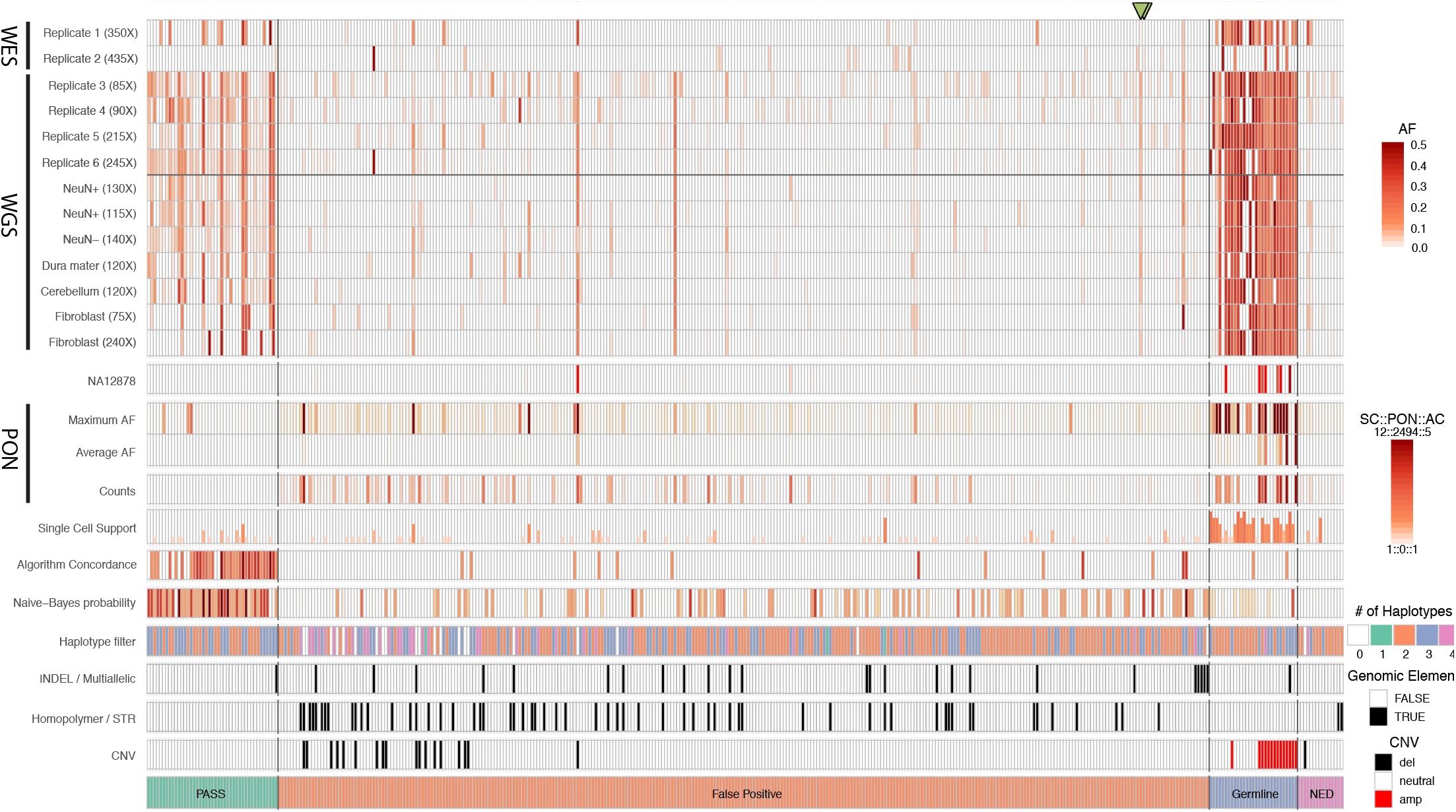
Summary of validation results for 400 candidate mosaic SNVs. Vertical lines represent candidate mosaic SNVs. Shaded rectangles to the right of the figure provide the keys to interpret the shading presented for each candidate SNV. There was concordance in true positive mosaic SNV calls (PASS; green rectangle at bottom of figure) in multiple datasets and secondary validation experiments. Chromium linked-read haplotype phasing and single cell sequencing datasets also were effective in supporting a subset of *bona fide* mosaic SNV calls. By comparison, the VAFs of falsepositive calls (red rectangle) are inconsistent across different datasets and often occur within or near insertion/deletion (indel) mutations, short tandem repeat sequences (STRs), homopolymeric nucleotide stretches, or copy number variants (CNVs). Importantly, the panel of normals (PON) filter, but not the comparison to WGS data from a control sample (*i.e.*, to NA12878), was highly effective at identifying contaminating false-positive SNV calls (orange rectangle) and germline SNPs (gray rectangle). We lacked sufficient data to evaluate a subset of candidate SNVs (purple rectangle, NED – not enough data). The two green triangles at the top of the figure denote mosaic SNVs that validation experiments deemed to be falsepositive calls; however, cell lineage analyses demonstrated that they are likely *bona fide* mosaic SNVs (see text & Fig. 4).

### Sequencing and/or mapping artifacts

The largest proportion of false-positive calls (300/312) were represented by fewer than five supporting reads (*i.e.*, low read count false-positive calls) in the 250X coverage replicate 6 WGS dataset, which had the highest coverage among all WGS datasets (Fig. 3). These false-positive calls likely represent DNA sequencing and/or mapping errors. Indeed, the use of Chromium 10X linked-read datasets allowed us to unambiguously identify SNV calls that resulted from DNA sequencing artifacts. Twenty-four false-positive calls appeared as alternative reads on both parental haplotypes, leading to the appearance of four apparent haplotype states (Fig. 3; see “4 haplotype” calls) and are most simply explained by sequencing errors in reads originating from both parental haplotypes. By comparison, *bona fide* validated mosaic SNVs were detected as alternative reads on a single haplotype, giving the appearance of three haplotype states (*i.e.*, the two parental haplotypes plus a “third” haplotype containing the somatic SNV on a single parental haplotype; **Fig. S6**).

The effectiveness of applying Chromium 10X linked-read datasets as a filter to evaluate *bona fide* somatic SNVs is limited by sequencing coverage. For example, at ~70X coverage, mosaic SNVs at <0.03 VAFs were unlikely to have supporting alternative reads and therefore could not be assigned to a parental haplotype (Fig. 3; see “2 haplotype” for validated SNVs). Similarly, the relatively small number of single cells analyzed in this study limited the effectiveness in using these data to validate mosaic SNVs present at low VAFs that were supported by independent PCR amplicon-based sequencing and/or ddPCR experiments (see below). Indeed, ~58% of validated and ~36% of false-positive SNVs lacked supporting alternative SNV reads in the single cell datasets.

### Genomic context

We noticed that candidate mosaic SNVs reads located near or within polymorphic germline indels, short tandem repeats, or structural variants (which includes copy number variants and retrotransposon insertions) were the second largest source of false-positive calls. A systematic misalignment of these germline variants (**Fig. S7**) and/or a systematic reduction in read counts near the germline variant (**Fig. S8**) resulted in 75 false-positive calls (**Table S2**). The systematic nature of these false-positive calls suggests they could be recurrently called as mosaic SNVs in unrelated samples. By extension, because sequencing errors and misalignments are known to be a problem in certain sequence contexts, some of the low read count false-positive calls and germline SNPs discussed above may also be due to systematic sequencing errors.

### Unrelated samples and replicate samples

We next considered a filtering strategy that is based upon analyzing read support for a given mosaic SNV call in unrelated individuals. We searched for alternate reads containing the 400 candidate mosaic SNVs using ~30X coverage WGS data from the 2504 unrelated samples in the 1000 Genomes Project (ftp://ftp-trace.ncbi.nlm.nih.gov/1000genomes/ftp/1000G2504highcoverage/). This analysis revealed that 183/400 calls were present at >0.05 VAFs in more than five individuals and likely represent systematic false-positive calls or germline variants. Indeed, 17/183 were germline, 153/183 were false positives, and 13/183 require addtional data to classify (*i.e.*, not enough data [NED] Fig. 3). This panel of normals (PON) filter provided discriminative power to detect false-positive calls that likely arise from systematic technical errors encountered during Illumina sequencing and/or read misalignments in certain genomic contexts across the VAF spectrum. For example, we found instances where sequencing errors occurred within short tandem repeats (*e.g.*, an A>T at chrX:150,725,216 in the sequence of 5′-CTC[A>T]CTCTCTCTCT; also see below). Given its importance, we calculated the PON mask across the entire human genome and have made these data available to the scientific community (https://bsmn.synapse.org).

Next, we searched for alternate reads in sequencing data derived from additional brain regions of the common reference sample. These datasets consisted of 75-240X WGS data derived from the cerebellum, neuronal and non-neuronal cell fractions of the dorsolateral prefrontal cortex (DLPFC), dura matter, and dural fibroblasts. Genotyping the 400 candidate variants using high stringency mapping quality (q30) and base quality (Q30) sequencing reads revealed consistent support for both germline SNPs and *bona fide* mosaic SNVs assignments across multiple datasets (Fig. 3). Namely, all the germline and 35/43 validated somatic SNVs were present at >0.01 VAFs in at least three replicate datasets, which is consistent with previous studies^17^. However, we observed that 9/12 false-positive calls with substantial read support (>=5) and 12/18 false-positive calls with a VAF >0.02 passed the filtering criteria, but likely arose from systematic sequencing errors. Thus, sequencing multiple brain regions and different tissues from the common reference sample is powered to confirm *bona fide* mosaic SNV calls, but does not replace the requirement of using the PON filter to remove systematic falsepositive calls.

### Best practices for calling mosaic SNVs

Consolidation of the above validation approaches led to the identification of 43 *bona fide* somatic SNVs. We next sought to derive a unified set of best practices to call somatic SNVs in WGS data. We found the input “discovery” call set has a significant impact on final sensitivity and determined that a GATK ploidy option of 50 for the 250X coverage WGS replicate 6 dataset, which corresponds to approximately five sequencing reads per haplotype, generally allowed the best approach to identify validated mosaic SNVs (Figs. 4d **and S9**). We applied a number of filters to remove false-positive calls (see below). Although the filters were effective, integrating a machine learning approach that unifies multiple read and genomic features^32^ into the best practices framework resulted in the highest accuracy call sets (**Fig. S9**).

**Figure 4:**
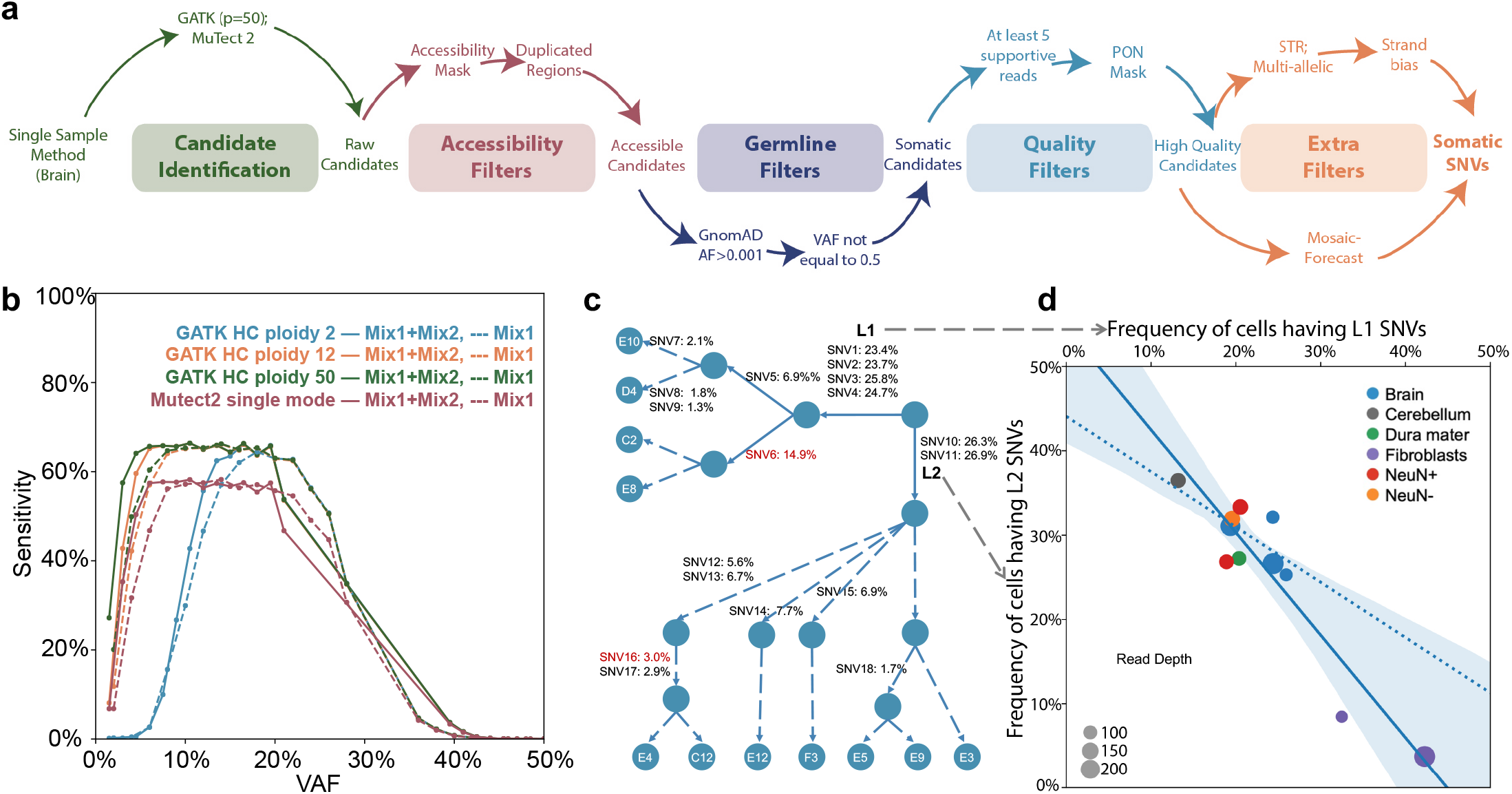
Best Practices Workflow to Call Mosaic Variants. (*a*) Schematic of the filtering strategies used to call mosaic SNVs using WGS and WES data. (*b*) GATK (at different ploidy settings) and Mutect2 were used to call simulated mosaic SNVs at different VAFs (x-axis) and sensitivities (y-axis) in dNa mixing experiments. (*c*) Reconstructed cell lineage trees using a cohort of mosaic SNVs (**Table S2**) present in eleven single-cell datasets from the common brain sample. Indicated are the names of each SNV (sNv1, SNv2,*etc*.) and the estimated SNV VAFs (from 250X WGS data). (*d*) SNVs marking the L1 (x-axis) and L2 (y-axis) lineages show anti-correlated VAFs across multiple brain and tissue samples, suggesting these SNVs differentiate the earliest cell lineages in this sample. Solid line, linear regression of the SNV anti-correlation values across all samples. Shaded areas is the corresponding 95% confidence intervals. Dashed line, linear regression of the SNV anti-correlation values when only brain samples are included in the analysis.

Based on the above data, the BSMN has the following recommendations for calling somatic SNVs from WGS data derived from non-cancerous cells (Fig. 4a, see **Methods**): (i) call variants with GATK using a ploidy setting that corresponds to 20% of the overall sequencing coverage (*e.g.*, a ploidy value of 50 for 250X WGS coverage); (ii) eliminate inaccessible genomic regions using the 1000 Genomes Strict Mask; (iii) discard germline variants that have a population allele frequency of >0.001 (*e.g.*, by comparing the candidate mosaic SNV calls to established catalogues of population variations such as the Genome Aggregation Database^33^; (iv) eliminate calls having ~0.5 VAFs by binomial test significance with a p-value cutoff above 10^−6^; (v) carefully scrutinize calls in genomic regions exhibiting copy number gains and other structural variants; (vi) mandate candidate mosaic SNVs have at least five independent non-duplicated supporting reads that have minimum values of 20 for mapping and base quality; (vii) identify and eliminate false-positive calls using the PON filter based on the data from the 1000 Genomes Project; (viii) filter calls using available machine-learning algorithms (*e.g.*, MosaicForecast^32^); (ix) optionally interrogate calls by applying filters that identify forward-reverse read imbalances and/or haplotype information (*e.g.*, 10X Genomics linked-read haplotype datasets); and (x) optionally validate candidate mosaic SNVs using additional wet-bench experimental approaches (*e.g.*, PCR amplicon-based sequencing and/or ddPCR). This best practices workflow is freely available at: https://github.com/bsmn/bsmn-pipeline.

Applying the first seven steps of this best practices workflow to 250X WGS replicate 6 data from the common DFPLC brain sample resulted in the identification of 40 candidate mosaic SNVs with >0.025 VAFs; 34 of these SNVs also were present in our original validation set. Two additional SNVs, which are absent from the set of 43 validated mosaic SNVs, also are likely true variants (discussed below). As an additional benchmarking metric, we used the best practices workflow to detect simulated SNVs in the DNA mixing experiment; however, because the simulated mosaic variants actually represent germline SNPs, we omitted the PON filter for this analysis. The sensitivity to detect simulated SNVs across the entire genome at VAFs ranging from ~0.02 to ~0.25 was ~65% (Fig. 4b), but increased to ~90% when we only considered simulated SNVs present in the accessible portion of the genome. The latter estimate is comparable to what we observed for a subset of 35 validated mosaic SNVs with >0.02 VAF (34 validated out of 35 [~97%]).

### Using somatic SNVs to recreate cell lineage trees

We next asked whether the identified mosaic SNVs could be used to infer developmental cell lineage trees. We first examined whether the union set of 49 SNVs identified in our validation call set and by the best practices workflow were shared among the twelve NeuN+ neuron single cell datasets. One neuron (**Fig. S10**, cell E2) was excluded from this analysis, as it lacked evidence for any of the 49 SNVs, perhaps because of inefficient whole genome DNA amplification. The VAFs of the 49 SNVs also were estimated from the following WGS datasets: DLPFC (4 samples), NeuN+ (2 datasets) and NeuN-(1 dataset) DLPFC cell fractions, cerebellum (1 dataset), dura mater (1 dataset), and dural fibroblasts (2 datasets). Hierarchical clustering using the single cell genotype and SNV vAf data allowed the reconstruction of a cell lineage tree (Fig. 4c; see **Methods**). Two lineages (L1 and L2) were clearly separable by diagnostic mosaic SNVs in the eleven single cells (Figs. 4c and **S10**; compare SNVs 1-4, which define the L1 lineage, with SNV10 and SNV11, which define the L2 lineage). Moreover, a subset of SNVs could be used to infer L1 (*e.g.*, SNV5, SNV6, SNV7, *etc*.) or L2 (*e.g.*, SNV12, SNV14, SNV18, *etc*.) sub-lineages. Notably, the L1 and L2 lineages were evident in ectodermally-derived brain tissue, ectodermally- and mesodermally-derived dura mater, and mesodermally-derived dural fibroblasts, suggesting the L1 and L2 lineages originated prior to gastrulation and diverged during early embryonic development, possibly after the first post-zygotic cleavage.

We next tested whether we could discriminate the presumed sister L1 and L2 lineages without relying upon the use of single cell sequencing data. We reasoned that frequencies of SNVs defining the L1 and L2 lineages would covary across DNA samples derived from different tissues of the common reference sample. For example, if the fraction of cells from the L1 lineage predominates in one tissue sample, we should observe a corresponding decrease in the fraction of cells from the L2 lineage in that tissue sample and *vice versa*. Moreover, the sum of allelic frequencies of SNVs defining the L1 and L2 lineages should approach 0.5 VAF (which corresponds to total cell frequency of 100%). We compared the VAFs of all 49 mosaic SNV pairs in nine brain region and two dural fibroblast samples sequenced to at least 75X coverage and then calculated how well the VAF sums of each SNV pair matches 0.5 across the WGS bulk samples (see **Methods**). The eight best scoring SNV pairs consisted of a SNV (SNVs 1-4) from the L1 lineage and a SNV from L2 lineage (SNV10 and SNV11; **Fig. S11**). Although the VAFs of SNVs defining the L1 and L2 lineages differed among tissue samples, their sums consistently approached 0.5, which correspond to a cell fraction value of ~100% (Figs. 4c and 4d). Using a similar strategy, we also were able to identify anti-correlated mosaic SNV VAFs that defined L1 and L2 sublineages that likely arose later in development (**Fig. S11**). Thus, this analysis suggested the possibility of discovering and analyzing mosaic SNVs across multiple bulk tissues to reconstruct early developmental cell lineages.

The recreation of cell lineage trees served as an independent means to gain support for *bona fide* mosaic SNV calls. For example, a somatic SNV defining a L1 lineage sub-branch (SNV6: chr7:112,461,481 G>T) was not present in our call set of 43 validated SNVs (even though it “passed” the Chromium 10X linked-read haplotype test) because it had a very low VAF (<0.0002) in PCR amplicon-based sequencing experiments. However, SNV6 exhibited a consistent vAf across multiple brain datasets and no apparent alignment or sequencing artifacts surround the site. In addition, and consistent with the reconstructed lineage tree (**Figs. 4c & S10**), the sum of the SNV5 and SNV6 VAFs for L1 sub-lineages was in general agreement with the VAF of the SNVs defining the L1 lineage across samples. Based on these analyses, SNV6 is likely a *bona fide* mosaic SNV. Using a similar strategy, we also concluded that a mosaic SNV defining a L2 sub-lineage (SNV16: chr6: 96,086,198 A>C) that was not in our original validation set also is likely is*a bona* fide somatic SNV (see **Methods**). Thus, in total, we were able to validate 45 somatic mosaic SNVs in the common reference sample.

## Discussion

The BSMN has described the successful implementation of a coordinated, multi-institutional approach to discover somatic SNVs in a neurotypical brain sample. These efforts led to the discovery of a catalog of high-confidence mosaic SNVs that are present at >0.01 VAFs at ~65% detection sensitivity; 42/43 validated somatic SNVs present at >0.02 VAFs in at least one WGS replicate. When adjusted for discovery sensitivity, this number of somatic SNVs is consistent with previous estimates that ~1.3 SNVs arise per cell division per cell during early development^17^.

A major factor currently affecting the sensitivity of our approach (towards discovering mosaic variants at a >0.02 VAF) is the necessity of restricting variant calling to the portion of the genome that can be assessed using short-read DNA sequencing technologies. The refinement of single-molecule long-read sequencing approaches (*e.g.*, Pacific Biosciences and Oxford Nanopore technologies) should enable SNV calling in currently inaccessible genomic regions; however, the sequence depth required for mosaic SNV discovery currently make the application of these approaches prohibitively expensive.

The use of orthogonal approaches to interrogate somatic SNV calls was instrumental in our validation analyses. In general, these approaches (*e.g.*, PCR amplicon-based sequencing, Chromium 10X-linked read sequencing, the analysis of single cell sequencing datasets, and ddPCR) were effective at identifying falsepositive calls and providing validation support for *bona fide* somatic SNVs, but likely are not suitable for genome wide *de novo* mosaic SNVs discovery. Moreover, each approach has limitations. For example, PCR amplicon-based sequencing initially led to the mis-assignment of a small number of *bona fide* somatic SNV calls as false positives. Similarly, the Chromium 10X linked-read and single cell sequencing data filters generally were only effective in supporting a small number of somatic SNV calls present at higher VAFs. Finally, although ddPCR proved to be a highly reliable validation approach, the effort and cost required for developing robust assays for each individual predicted mosaic SNV only allowed it to be applied to assess a small number of variants. It is noteworthy that distinguishing somatic SNVs present at low VAFs from false-positive calls (*e.g.*, sequencing artifacts or calls arising from read misalignment errors) remains a formidable challenge.

Our analysis provided tantalizing suggestive evidence that the use of somatic SNVs discovered in bulk WGS sequencing data may ultimately provide a cost-effective approach to reconstruct early embryonic cell lineage trees when compared to more expensive single cell sequencing and clonal analysis approaches. However, because we only analyzed a limited number of bulk DNA samples from a single individual, additional data will be needed to explore the potential advantages and limitations of tracing cell lineages from mosaic SNVs discovered in bulk WGS datasets.

In sum, the BSMN has developed an efficient approach to identify and validate somatic SNVs in the brain of a single neurotypical individual that should serve as a guide to ultimately assess the contributions of somatic SNVs to neuropsychiatric diseases. This study generated ~7 Tb genomic sequencing data that have been deposited into the NIMH Data Archive, which is freely available to the greater research community.

## Acknowledgements

Members of the BSMN consortiums are listed in the supplementary file. We thank Geetha Senthil, Miri Gitik, and Thomas Lehner for their help organizing the BSMN consortium and promoting the study. We express our gratitude to the NDA team for providing collaborative data storage space and support for data transfer needed for the successful completion of this study. This study was supported by NIMH grants U01MH106892, U01MH106876, U01MH106883, U01MH106874, U01MH106893, U01MH106882, U01MH108898, U01MH106891 and U01MH106884.

## Data availability

Data and call sets have been deposited in the NIMH Data Archive (NDA Study ID 792: https://dx.doi.org/10.15154/1504248) and can be accessed as part of the NIMH Data Archive permission groups: https://nda.nih.gov/user/dashboard/datapermissions.html. The PON mask applied in the study can be accessed from: https://www.synapse.org/#!Synapse:syn22024464.

## Methods

### Data generation

#### Mixing Experiment

Lymphoblastoid cell lines (LCLs) of four grandparents from CEPH/Utah pedigree 1463 were ordered from the Coriell Institute (catalog numbers: GM12889, GM12890, GM12891, GM12892). Genomic DNAs (gDNA) from the cells were prepared, mixed in different proportions (as indicated in Fig. 1a), and subjected to WGS. Briefly, DNA libraries were prepared from 1μg of mixed gDNA per sample using the Illumina PCR-free TruSeq DNA Library Prep kit according to the protocol provided by the manufacturer. The constructed libraries were quantified using the KAPA Library Quantification Kit and real-time PCR. Then, 150 bp paired-end reads were generated using the Illumina HiSeq X platform. The sequencing experiments were designed to yield ~200X coverage on each samples and were carried out at a GeneWizM facility (South Plainfield, NJ).

#### Sample processing (*Lieber Institute, contributed by Daniel Weinberger*)

The samples from the common reference brain numbered 5154 were dissected from deep frozen tissue from various brain regions in a standardized routine that uses a dental drill to minimize tissue injury and RNA degradation. To ensure that all distributed samples came from a singular source, the brain samples were first uniformly homogenized and then aliquoted for distribution to all but one BSMN node (Yale University and Institut de Biologia Evolutiva), which received a piece of frozen tissue. The disbursed tissue and related data were deidentified, delinked, and coded prior to sharing with the BSMN nodes. To obtain viable fibroblasts we collected a biopsy from the meninges (dura mater) of the same individual, suspended it in appropriate media (including DMEM (Gibco), HiFBS (Gibco), GlutaMAX (Gibco), Antibiotic/Antimycotic Solution (Gibco), MEM NEAA (Gibco), 2 βME (Gibco)), and then cultured it into a fibroblast line. The resultant fibroblast cell lines were serially expanded in T-25 and T-75 tissue culture flasks, harvested, re-suspended in media as a large homogenous culture, and subsequently were aliquoted into smaller tubes for storage/redistribution.

#### Whole Exome Sequencing Replicate 01 (*University of Michigan, contributed by Sarah Emery and Yifan Wang*)

Genomic DNA was extracted from 5154 brain tissue using the MagAttract HMW DNA Kit (Qiagen, Germantown, MD). The length of gDNA was determined by either standard gel electrophoresis in 0.4% agarose or pulse field gel electrophoresis in 1% agarose and 0.5 × TBE for 16 hours at 6 V/cm and a 1200 angle with an initial switch time of one second and a final switch time of six seconds. Whole exome sequencing was performed on all extracted gDNAs. Duplicate libraries were made for each sample by shearing 75-200ng of gDNA to 350bp. Libraries were purified with 0.65x SPRIselect beads (Beckman Coulter) and quantitated using a Qubit™ dsDNA HS Assay Kit (Thermo Fisher Scientific, Carlsbad, CA). A 50ng aliquot of sheared DNA was saved for later use and the remaining 400-800ng was used for exome target enrichment. Target enrichment was performed using SeqCap EZ Exome Probes v3.0 (Roche Sequencing Solutions, Pleasanton, CA) according to the protocol provided by the manufacturer with a 72-hour incubation for hybridization and 12-16 cycles of post-capture ligation-mediated PCR (LM-PCR) to amplify the captured DNA. The quantity of the captured DNA was measured using a Qubit™ dsDNA HS Assay Kit and target enrichment was determined by calculating the abundance of control targets in post-capture libraries relative to the abundance of these targets in the pre-capture libraries as outlined in SeqCap_EZ_UGuide_v5.4 (Roche Sequencing Solutions). Each library was sequenced on an individual HiSeq lane on the HiSeq X series. Library QC and sequencing was performed at Novogene Corporation (Davis, CA).

#### Whole Exome Sequencing Replicate 02 (*University of California San Diego, contributed by Laurel Ball*)

Pulverized brain cortex (0.98g) and fibroblasts were provided by the Lieber Institute for Brain Development, (Baltimore, MD). Fibroblasts were cultured in fibroblast culture media containing MEM (Gibco), 20% FBS (Gibco), and 1X Pen/Strep (Gibco). Confluent fibroblasts were harvested and gDNA was extracted from pulverized brain and fibroblast samples using Qiagen Maxiprep kits according to the protocols provided by the manufacturer. Genomic DNA samples were prepared for whole exome sequencing using the Agilent SureSelect XT Human All Exon v.5 kit. Then, 125 bp paired-end reads (median insert size ~210 bp) were generated using the Illumina HiSeq X 2500 platform. The sequencing experiments were designed to yield three datasets of ~100X coverage on each sample, sequencing completed at the New York Genome Center, NY.

#### Whole Genome Sequencing Replicate 03 (*Yale University and Institut de Biologia Evolutiva, contributed by Irene Lobon*)

A frozen piece of DLPFC from the 5154 common reference brain sample was provided by the Lieber Institute for Brain Development (Baltimore, MD). A scalpel was used to scrape off its surface and subdivide it into smaller pieces. Genomic DNA was extracted using the DNeasy Blood and Tissue kit (QIAGEN). The NEBNext Ultra II DNA Library Prep Kit for Illumina was used for library preparation. Then, 125 bp paired-end reads were generated using the Illumina HiSeq4000 platform. The sequencing experiments were designed to yield 90X coverage. Sequencing was carried out by Macrogen.

#### Whole Genome Sequencing Replicate 04 (*Lieber Institute University, contributed by Rujuta Narurkar and Joo Heon Shin*)

Nuclei were isolated from pulverized postmortem brain tissue (0.7g) using sucrose gradient centrifugation (4°C, 25 000 rpm, 1h) and incubated with anti-NeuN-488 antibody (Millipore MAB377X, 1:1000) in a phosphate buffered saline (PBS) solution containing bovine serum albumin (BSA at 0.1%). Nuclei were sorted into NeuN+ and NeuN-fractions by FACS (BD FACSAria™III). Fibroblasts were cultured in fibroblast culture media (DMEM high glucose, FBS, L-glutamine, N.E. amino acids, Pen/Strep; all Invitrogen) and confluent fibroblasts were harvested for gDNA isolation. DNA was extracted from: (i) 0.3g post mortem bulk brain tissue; (ii) NeuN+ and NeuN-nuclear fractions; and (iii) fibroblast cell lines using the DNeasy Blood and Tissue Kit (QIAGEN) that included an RNAse A treatment step (QIAGEN) according to protocols provided by the manufacturer. For all samples, the libraries were prepared using the Illumina TruSeq DNA PCR-Free Library Prep Protocol with 1.5μg of DNA for NeuN+ and NeuN-nuclear fractions and 3 μg of DNA for bulk cortex and fibroblast samples. Then, 150 bp paired-end reads were generated using the Illumina HiSeq X platform. The sequencing experiments were designed to yield 90X coverage. Library construction and sequencing were conducted at Psomagen, previously known as Macrogen (Rockville, MD, USA).

#### Whole Genome Sequencing Replicate 05 (*Harvard University, contributed by Javier Ganz*)

Pulverized brain tissue was provided by Lieber Institute for Brain Development (Baltimore, MD). DNA was isolated from bulk brain tissue using lysis buffer from the QIAamp DNA Mini kit (Qiagen) followed by phenol chloroform extraction and isopropanol cleanup. DNA was then processed at Macrogen using the TruSeq DNA PCR-Free library preparation (Illumina) followed by a minimum of 30X sequencing of seven separate libraries on the Illumina HiSeq X Ten, for a total minimum coverage of 210X.

#### Whole Genome Sequencing Replicate 06 (*Yale University, contributed by Liana Fasching*)

Pulverized brain cortex (1.02g) and fibroblasts were provided by the Lieber Institute for Brain Development (Baltimore, MD). Nuclei were isolated from pulverized postmortem brain tissue (0.7g) using sucrose gradient centrifugation (4°C, 25 000 rpm, 1h) and incubated with anti-NeuN-488 antibody (Millipore MAB377X, 1:1000) in PBS and BSA. Nuclei were sorted into NeuN+ and NeuN-fractions by FACS (Bd FACSAria™ III). Fibroblasts were cultured in fibroblast culture media (DMEM high glucose, FBS, L-glutamine, N.E. amino acids, Pen/Strep; all Invitrogen) and confluent fibroblasts were harvested for gDNA isolation. DNA was extracted from: (i) 0.3g post mortem bulk brain tissue; (ii) NeuN+ and NeuN-nuclear fractions; and (iii) fibroblast cell lines using the DNeasy Blood and Tissue Kit (QIAGEN) that included an RNAse A treatment step (QIAGEN) according to protocols provided by the manufacturer. Illumina Truseq DNA PCR-free (350bp insert) libraries were prepared for all samples using 1.5 μg of DNA for NeuN+ and NeuN-nuclear fractions and 3 μg of DNA for bulk cortex and fibroblasts. Then, 150bp paired-end reads were generated using the Illumina platform. The experiments were designed to yield 30X f sequence coverage for NeuN+ and NeuN-nuclear fractions and 210X sequence coverage for bulk cortex and fibroblast samples. Library construction and sequencing were conducted at Macrogen (Rockville, MD, USA).

#### Whole Genome Sequencing of single cells (*Yale University, contributed by Liana Fasching, Livia Tomasini and Bo Zhou*)

Nuclei were isolated from fresh frozen pulverized postmortem brain tissue using sucrose gradient centrifugation (4°C, 25 000 rpm, 1h) and incubated with anti-NeuN-488 antibody (Millipore MAB377X, 1:1000) in PBS and BSA. Ninety-five NeuN+ single nuclei were sorted into a 96-well plate by FACS (BD FACSAria™ III). Whole genome amplification was performed by multiple displacement amplification (MDA) using the REPLI-g Single Cell Kit (QIAGEN) according to protocols provided by the manufacturer. After amplification, each genomic DNA was purified using the DNeasy Blood Tissue Kit (QIAGEN). Quality control was performed using a multiplex PCR for four arbitrary loci from different human chromosomes^12^. Amplified single nuclei were subsequently excluded if less than four loci were amplified. Sixty seven out of ninety-five amplified nuclei (70.5%) passed the 4-locus multiplex PCR quality control. Low coverage sequencing was performed as a second quality control to assess locus dropout rate. Paired-end barcoded WGS libraries were prepared for the 67 MDA reactions and sequenced on one lane of HiSeq 4000 (2X100 bp). Twelve out of the sixty-seven MDA reactions (18%) were selected for further sequencing. Illumina Truseq DNA PCR-free libraries were prepared for seven of them and sequence on a HiSeq X (2X150 bp) at 30X coverage.

Samples of MDA amplified DNA from five cells was size selected for fragments >10 kb on the BluePippin instrument (Sage Science, Beverly, MA, USA) using the S1 Maker 10 kb High Pass protocol provided by the manufacturer and then diluted to 1 ng/μl and used as input for the Chromium reagent delivery system^34,35^ from 10x Genomics (Pleasanton, CA, USA), where high molecular weight (HMW) DNA fragments are partitioned into >1 million droplets, uniquely barcoded (16 bp) within each droplet, and subjected to random priming and isothermal amplification following standard manufacturer’s protocol. Afterwards, the emulsion was broken, and the barcoded DNA molecules were released and converted to a Chromium 10X linked-read library in which each library molecule retains its “HMW fragment barcode”. Read-pairs generated in this manner (*i.e.*, linked-reads) that come from the same HMW DNA fragment can be identified by their “HMW fragment barcode” and subsequently used to phase variants onto megabase scale haplotypes. The final libraries (8 cycles of PCR amplification) were diluted to 5 nM and sent to Macrogen (Rockville, MD, USA) for sequencing (2×151 bp) on the Illumina HiSeq X.

#### Chromium Linked-Read sequencing (*University of Michigan, contributed by Sarah Emery and Yifan Wang*)

Pulverized, frozen brain tissue and dural fibroblasts (FIBRO) from a deceased, neurotypical, male individual (5154) were received from the Lieber Institute for Brain Development (Baltimore, MD). Dural fibroblast were cultured in DMEM (Gibco/ Life Technologies) supplemented with 10% FBS (Gibco/Life Technologies), 2% Glutamax (Gibco/ Life Technologies), and 1% Antibiotic, Antimycotic (Gibco/Life Technologies). Cells were cultured for 3-10 weeks and passaged when they reached 85%-95% confluence.

Genomic DNA (gDNA) was extracted from 5154 brain tissue and dural fibroblasts using the MagAttract HMW DNA Kit (Qiagen, Germantown, MD). The length of gDNA was determined by either standard gel electrophoresis in 0.4% agarose or pulse field gel electrophoresis in 1% agarose and 0.5 × TBE for 16 hours at 6 V/cm and a 1200 angle with an initial switch time 1 second and a final switch time-6 seconds. For 5154 brain gDNA, 5154 FIBRO, an aliquot containing 1-5 μg of gDNA was sent to HudsonAlpha Discovery (Huntsville, AL) for linked-read sequencing using 10x Genomics technology (Pleasanton, CA). Long Ranger v2.2 (10x Genomics) was used to align reads and then call and phase SNPs to obtain haplotype information for each read.

#### Data uniform processing at Amazon Web Services (*Mayo Institute, contributed by Taejeong Bae*)

All sequencing data generated by each BSMN node were uploaded to the shared S3 bucket of the Amazon Web Services (AWS). We uniformly processed the data in the AWS system to prepare aligned bam files. We implemented a mapping workflow similar to the best practices protocol suggested by GATK. The original FASTQ files, or those obtained from the conversion of BAM files, were split by flowcell lanes using an in-house awk script so that reads in each lane form the same read group in alignment. For each sample, the reads from FASTQ files were aligned to the human reference genome GRCh37d5 (ftp://ftp-trace.ncbi.nih.gov/1000genomes/ftp/technical/reference/phase2referenceassemblysequence/hs37d5.fa.gz) using bwa (version 3.7.16a), sorted per each read group, and merged into a single BAM file with sambamba (version 0.6.7). The merged BAM files were marked for duplicate reads using PICARD (v2.12.1). Then, we performed indel realignment and base quality recalibration using GATK (v3.7-0), resulting in the final uniformed processed BAM files. This mapping workflow was installed and run in an AWS cluster prepared by AWS ParallelCluster (https://github.com/aws/aws-parallelcluster).

### Calling variants in the reference brain

#### Method 1 (*University of California in San Diego, contributed by Xiaoxu Yang*)

Both tissue-specific and tissue-shared mosaic variants were called from the WES sequencing data. Brain- and fibroblast-specific variants were called using Mutect2 (GATK3.8)^25^ and Strelka2^27^; the bam files from the brain sample (combined and non-combined from independent sequencing libraries) and fibroblast sample were treated as “tumor-normal” and “normal-tumor” pairs separately and cross-compared between each other. Variants called by both callers were listed. Mosaic variants shared between the brain and fibroblast samples were called using the single mode of MosaicHunter^36^ by either combing all brain replicates or calling each separate sample. Variants that passed all the MosaicHunter filters also were listed. Mosaic candidates from the combined lists were further filtered using the following criteria: (i) the variant had more than 3 reads for the alternative allele; (ii) the variant was not present in UCSC repeat masker or segmental duplications; (iii) the variant was at least 2bp away from a homopolymeric tract; and (iv) the variant exhibited a gnomAD allele frequency lower than 0.05. Variants that exist in the 1000 genome project (phase 3) also were excluded from the analysis. Variants from both exome data sources were tested and a combination of tissue-specific mosaic variants and tissue-shared mosaic variants were collected and the credible interval of VAFs were calculated using a Bayesian based method described previously^37^.

#### Method 2 (*Yale University and Institut de Biologia Evolutiva, contributed by Irene Lobon*)

Somatic SNVs were called for each single sample using VarScan 2^38^ with relaxed parameters (-min-coverage 1 -min-reads2 1 -p-value 1 and -min-var-freq 0.000001) to increase sensitivity. Calls were excluded if they overlapped the following somatic non-callable regions: (i) the 1000 Genomes strict mask non-pass positions; (ii) the WGAC segmental duplications track; (iii) the dbSNP common database; or (iv) positions with a mappability <1 for the corresponding read length. A Fisher exact test was used to filter out false-positive calls caused by random sequencing errors and a binomial test across the samples was used to remove germline heterozygous variants. Regions with higher coverage were determined using a Chi-squared test. Strand biased candidates were identified with a Poisson and a Fisher exact test. Variants within 5 bp of indels, near homopolymers, or within 300 bp of each other were also filtered during the analysis. Similarly, calls with a biased position in the reads for one of the alleles were also excluded from the analysis. Additionally, somatic SNVs were called with GATK 3.8^25^ Haplotype Caller (-ploidy 4) and filtered with the somatic non-callable, sequencing error, and binomial tests. The final call set consisted of mosaic SNVs passing all VarScan2 filters in at least three samples and all Haplotype Caller filters in at least one sample.

#### Method 3 (*Harvard University, contributed by Alon Galor and Max Sherman*)

To generate a list of high-confidence brain somatic variants for the common experiment sample, we considered three high-quality, high-depth Mixed-Brain samples generated from the Replicate 6, Replicate 4, and Replicate 3 datasets. We produced putative somatic variant calls for each of these three samples with Mutect2 (v2.3.5)^26^, using the highest-depth common experiment fibroblast sample available, generated by Replicate 6, as a matched normal sample. Only variants flagged as “PASS” were retained for further analysis. We then leveraged Mutect2 to generate a Panel of Normals, using 75 200X WGS Brain samples. To construct this panel, we ran Mutect2 in its tumor-only mode on each of these 75 samples, making use of non-default parameters to produce more lenient calls. If a putative call from the common experiment sample was present at greater than a 1 % allele fraction in two or more of the Panel of Normals samples, it was filtered out. Subsequently, we removed putative calls coinciding with Mutect2 tumor-only calls (again, using the same non-default parameters described above) of the remaining Mixed-Brain fibroblast samples. Next, we excluded putative somatic variants in the 1000G, ExAC, and ESP5600 common variant databases, as well as those located in inaccessible regions, according to criteria outlined by the 1000 Genome Project Strict Mask. We also removed putative somatic variants in indel, copy number variant, segmental duplication, and structural variation genomic regions. To generate indel calls, we employed the GATK Best Practices Germline Variant Calling Workflow. Variants in structural variant regions were defined as those in a 200bp region, centered at the SNV, where the number of clipped reads was in the ≥ 95th percentile of 200bp regions genome-wide. Variants were deemed to lie in a copy number variant or segmental duplication region if a 200bp region, centered at the SNV, had GC-normalized depth in the ≥ 95th percentile of 200bp regions genome-wide.

In order to prioritize variants, we further tiered them using evidence available for the calls in additional bulk brain tissue samples and single brain cells. In particular, we annotated putative SNVs observed on exactly one haplotype in phased bulk tissue, deemed significantly present in other Mixed-Brain samples by a joint-genotyping likelihood method, or present in single cells, as determined by an in-house method for analyzing single cells for evidence of mosaic events.

#### Method 4 (*Kennedy Krieger Institute, contributed by Jeremy Thorpe*)

Candidate somatic variants were identified leveraging high-depth whole genome sequencing of four brain prefrontal cortex and cultured dural fibroblast samples. Paired brain-fibroblast variant calling was performed with MuTect2 (v4.0.1.0)^26^, Strelka2 (v2.8.4)^27^, and MosaicHunter (v1.0)^36^. Additionally, single brain samples were analyzed with the GATK (v3.8-0) HaplotypeCaller with ploidy 5 parameter. Germline InDels and CNVs were called with the GATK Best Practices Germline Variant Calling Workflow and CNVkit (v1.1), respectively.

Quality filters were applied to mosaic SNV candidates. Variants were excluded if they: (1) occurred in error-prone genomic intervals, as defined by the 1000 Genomes Strict Mask and/or UCSC RepeatMasker; (2) clustered within 1 kbp; (3) were within the 50 bp adjacent to InDels; (4) had a read depth >2 standard deviations of mean coverage; (5) were likely germline events occurring as GnomAD single nucleotide polymorphisms >0.1% allele frequency; (6) occurred within a CNV; (7) exhibited a read strand bias using binomial test; (8) were heterozygous sites using binomial test; (9) failed a Fisher’s exact test for allele strand bias; (10) had minor allele frequency <0.01 to exclude likely false-positive, low frequency events. Following removal of low quality sites, candidates were tiered using additional bulk and single cell datasets.

We tiered candidates by leveraging biological replicates, bulk 10X sequencing, and single cell data. Candidates were tiered on supporting evidence of the mosaic allele in multiple independent replicates, single cell allele support, and concordance of mosaic alleles within a single haplotype by haplotype phasing of 10X data.

#### Method 5 (*Mayo Clinic, contributed by Taejeong Bae*)

For calling somatic variants, we applied a single sample approach using the GATK (3.7-0) Haplotype Caller to each high-depth WGS dataset. To increase sensitivity towards calling somatic variants present at <10% variant allele frequencies, we used ploidy options of 2 to 10 for GATK Haplotype Caller. We took only “PASS” calls from each raw call set. Those “PASS” SNVs were filtered out using common variant databases, such as the 1000 Genomes, ExAC, GnomAD, ESP5600, and Kaviar to exclude known germline variants. Then, we filtered out the SNV sites residing in the genomic region of non-P bases of the 1000 Genomes Project Strict Accessibility Mask to filter out calls in problematic genomic regions. Remaining heterozygous germline variants were excluded by applying binomial test (P<0.00001), which identifies variants with 0.50 VAFs given the numbers of total reads and supporting reads. For checking strand bias of each candidate site, we used binomial test (P<0.05) for evenness of the counts of both strand reads and a Fisher’s Exact test (P<0.05) to identify imbalance of strand ratios between reference (REF) and alternative (ALT) bases. We filtered out the sites having either bias. We excluded the sites having more than two alleles (multi-allelic sites). Finally, the germline CNV status for the +/− 1kb region of each candidate site was checked using the genotype function of CNVnator. We removed the sites with >2.5 estimated copy number as likely duplicated regions.

#### Method 6 (*University of Michigan, contributed by Yifan Wang*)

Candidate variants from paired brain and dural fibroblast samples were called using MuTect and Strelka with the default parameters. At the same time, candidate variants from single brain samples were called using the GATK Haplotype Caller with parameter of ploidy set to 5. Candidate mosaic SNV sites then were filtered out using multiple quality filters. We first filtered out variants that overlapped with repetitive regions or low mappability regions, including regions covered by UCSC RepeatMasker simple repeats, Segmental Duplications, Simple Repeat tracks, regions not covered by 1000 Genomes Project Strict Accessibility Mask, as well as any variants not within +/− 3 standard deviations of mean sequencing coverage. We excluded common variants in GnomAD with a population allele frequency larger than 0.1%. During the process of counting alleles at candidate positions, reads with mapping quality scores lower than the 90 percentile of control sites (sites with high confidence), with more than 3% mismatches, as well as candidate sites exhibiting a base quality score lower than 20 were excluded from further analysis. After accounting for different alleles at candidate positions, we excluded sites with a candidate allele frequency larger than 0.01 in the control sample NA12878 (from the Genome in a Bottle Project). We applied a Fisher’s Exact test to exclude the sites whose alternative alleles are enriched on one strand compared to the other. We then filtered out sites with a known indel within +/− 5 base pairs of the variant. In the end, we had an allele frequency cutoff at 0.03 to exclude the extremely low frequency sites.

After removing the low quality sites, we applied a binomial test with false-discovery protection using the Benjamini-Hochberg procedure^39^ to filter out the heterozygous sites. We also used the haplotype information from the 10X linked-read common reference brain sequencing dataset to further filter out false-positive sites. We additionally removed candidate sites located within 100 bp of each other.

### Validation with targeted PCR amplicon-based sequencing approaches

Different PCR amplicon-based sequencing approaches were used to validate mosaic SNV candidates identified using the different computational pipelines. Each of the validation procedures is described below.

#### Amplicon 1 (*Yale University, contributed by Liana Fasching and Simone Tomasi*)

We validated the called SNVs by PCR amplicon-seq using following samples: (i) reference tissue – pulverized cortex DNA; (ii) reference tissue – fibroblast DNA; and (iii) control lymphoblastoid cell line DNA – NA12878. We determined the genomic location of each SNV using the Genome Browser. We designed each primer pair by selecting an 800 nucleotide long DNA template surrounding the SNV (400 nucleotide up- and 399 nucleotides down-stream of each SNV) using Primer3^40^. The specificity of primers was confirmed by UCSC *in silico* PCR (http://genome.ucsc.edu/cgi-bin/hgPcr) (**Table S5**).

The PCR amplification was performed using the Phusion High-Fidelity DNA polymerase (ThermoFisher). The optimal annealing temperature was predicted using the Tm calculator tool provided by the ThermoFisher website. The presence and size of each amplicon was confirmed by gel electrophoresis (2% agarose gel). In samples where we detected multiple PCR products, DNA bands with the correct amplicon sizes were extracted via gel extraction and purification using 2% agarose precast gels (E-Gel EX, Invitrogen) and gel imager (E-Gel Safe Imager connected to E-Gel iBase, Invitrogen), according to the recommendations provided by the manufacturer. Amplified DNA fragments were purified using the QIAquick PCR Purification Kit (Qiagen). Samples were multiplexed and sequenced under following conditions: MiSeq paired-end, 300 bp, 3 samples/lane.

#### Amplicon 2 (*University of Michigan, contributed by Sarah Emery and Yifan Wang*)

Putative mosaic SNVs were validated by high throughput sequencing of amplicons that contain the SNV and then calculating the relative abundance of reads containing the alternate allele. Primers were designed using Primer 3 software^40^ with 300-400 bp of genomic sequencing surrounding the SNV as the input. Since the read length of the amplification product is ~300 bp, we were able to gain an overlapped region between the paired reads. With the overlapped regions containing the somatic SNV candidates, we were able to sequence each candidate site twice and increase the accuracy of sequencing result by excluding reads with non-concordant bases from the pair ended reads at the candidate SNV positions. If possible, SNPs known to be heterozygous in our samples were included in the amplified sequence used as input. Primers were tested in silico^41^ to confirm they uniquely target the correct region of the genome. Phusion^®^ High-Fidelity DNA Polymerase (New England Biolabs) was used according to instructions provided by the manufacturer for amplification and primers were cycled under varying conditions to determine optimal PCR mix and annealing temperature. To generate amplicons for sequencing, either NA12878 or gDNA from the 5154 common reference brain tissue was used as template. The PCR product was purified with 0.7x SPRIselect beads (Beckman Coulter) and 10% of product was visualized on an agarose gel to confirm that only one amplicon of the correct size was present. If the size of the amplicon agreed with the size predicted from the primer design, we then sequenced the targeted amplified genomic fragment using an Illumina MiSeq sequencer. If the size was incorrect, we designed a second set of primers to obtain unique amplification. If none of the primers worked, the candidate was flagged with ‘primer not designed,. Protocols and reagents from NEBNext^®^ Ultra™ DNA Library Prep Kit for Illumina^®^ (New England Biolabs) were used for end repair, dA-tailing, and to ligate NextFlex adapters (Perkin Elmer, Waltham, MA) onto amplicons. After ligation, reactions were purified with 0.7x SPRIselect beads (Beckman Coulter) and PCR enrichment of adapter-ligated DNA was performed for 10 cycles using NEBNext^®^ Ultra™ DNA Library Prep Kit (New England Biolabs). Amplified libraries were purified with 0.7x SPRIselect beads and sequenced with MiSeq Reagent Kit v3, 600 cycle PE on MiSeq sequencer (Illumina, San Diego, CA).

#### Amplicon 3 (*Mount Sinai School of Medicine, Chaggai Rosenbluh*)

We sent all 400 target sites to Paragon Genomics and obtained a CleanPlex Custom NGS Panel allowing us to perform two multiplex PCR-based targeted resequencing assays for a total of 382 target sites. Eighteen of 400 sites were deemed not reliably targetable. The panels were designed using a proprietary primer design algorithm (sequences are available upon request) and proprietary background cleaning and molecular barcoding technologies to amplify the regions of interest and attach adapters for Illumina sequencing. The panel was iteratively optimized in-silico to prior to amplifying DNA from the common sample. The amplicons ranged in size from 250 bp to 425 bp (including the adapter sequences). A single MiSeq run provided ample coverage for all successfully targeted sites.

#### Amplicon 4 (*Harvard University, contributed by Javier Ganz*)

Targeted validation was attempted on 100 candidate mosaic variants and primers for 98 of those were successfully generated using BatchPrimer3^42^. PCR primers were synthesized with flanking Ion Torrent adapters P and A. Each primer pair was unique and their sequence served as a barcode for variant identification. Phusion HotStart II DNA Polymerase (Thermo)CR was used for DNA amplification over 25 cycles following manufacturer guidelines. PCR products were pooled and purified using AMPure XP technology (Agencourt) to be sequenced on the Ion Torrent S5 platform (Ion 530 chip). After demultiplexing and trimming, reads were mapped using BWA-MEM and locally realigned with GATK. Average coverage of mappable reads was 96350X per variant. Mosaic fractions were identified using custom scripts based on SAMtools mpileup.

#### Amplicon 5 (*Kennedy Krieger Institute, contributed by Jeremy Thorpe*)

We validated candidate mosaic SNVs by targeted, high coverage amplicon sequencing. PCR primers were designed with Primer3^40^ to generate amplicons from 300-500 bp and *in silico* verified to minimize the possibility off-target products. Amplicon sizes and purity were confirmed by PCR followed by agarose gel electrophoresis. Amplicons were quantitated, purified, and pooled for each of the NA12878 and 5154 brain tissue samples. NGS libraries were prepared using Illumina TruSeq Nano DNA kit and following standard Illumina NGS library preparation protocols. Libraries were sequenced with MiSeq Reagent Kit v3, 600 cycle PE on a MiSeq. Following demultiplexing and trimming, paired-end reads were merged with FLASH^43^ and aligned with BWA-MEM to hs37d5 human reference genome. The average depth of sequencing was 103335x per candidate. Target mosaicism was assessed by SAMtools mpileup and custom scripts.

#### Amplicon sequencing data analysis pipeline (*contributed by Yifan Wang*)

We constructed a framework for uniform processing of pCr amplicon validation data. The PCR amplicon paired-end sequencing data was assembled to a single read using PEAR^44^ with the non-concordant bases between the two reads set to N with a base quality of zero. We then aligned the reads to GRCh37d5 using bwa mem. Indel realignment was performed using the Genome Analysis Toolkit^25^. After pre-processing, we applied a series of filters to evaluate the putative mosaic SNV calls. We required a minimum of 200 reads covering the candidate mosaic SNV prior to making a “call” decision. Sites with fewer than 200 reads covering the candidate SNV were assigned the flag of ‘read not enough,. We then compared the allele fractions of the candidate mosaic SNV in the amplicon data from common reference brain sample and from NA12878. Since the same mosaic SNV event is unlikely to occur in two different individuals, we reasoned that *bona fide* mosaic SNVs should only be present in the brain sample. We applied a hard cutoff that removed any site with a candidate allele fraction >0.01 in NA12878 and additionally applied a skellam test to compare the putative mosaic SNV allele frequencies in the two samples. Together, these criteria excluded likely false-positive mosaic SNV calls that arise from sequencing errors in certain genomic contexts. We additionally established an empirical error model to exclude false-positive mosaic SNVs that likely arose during DNA amplification, library preparation, and/or sequence processing. We further evaluated PCR amplicon sequencing errors by assessing the mismatch rate (second allele frequency) in the overlapping region of the paired-end reads. We then used the 95^th^ percentile of the mismatch rate distribution found for each type of base change as the sequencing error cutoff for base changes at the candidate positions. Finally, we excluded candidate mosaic SNVs with >0.40 VAFs, as they likely represent germline SNPs.

We performed some additional final checks on the validated mosaic SNV events. First, we tested whether there was a positive correlation between the putative mosaic SNV VAFs in the WGS and PCR amplicon-based sequencing experiments (**Fig. S12**). Second, we tested whether there was a positive correlation between the ratios of the second highest allele frequency and the third highest allele frequency with the putative somatic SNV VAF in the WGS dataset (**Fig. S12**); the third highest allele frequency did not correlate with the variant allele frequency (**Fig. S12**).

#### Variant validation using single cells (contributed by Alon Galor and Max Sherman)

To distinguish a true mosaic variant from a multiple displacement amplification artifact in single cell sequencing, we adapted the method of Dong et al.^24^. First, we augmented their kernel smoothing method to utilize phased genotypes^45^ as this improves estimation of allelic imbalance at a given locus. Phasing was performed on the GATK HaplotypeCaller germline variants from Replicate 6 bulk brain tissue using Eagle^46^ and the 1000 Genomes Phase 3 reference panel^47^. Second, we determined the optimal kernel size to maximize accuracy using PaSD-qc^48^, which was determined to be 10 kb. Third, we instituted a likelihood ratio test to distinguish a first-round amplification artefact from a true mosaic variant; let *f(X;p*) be the probability of observing reads *X* where *p* is the expected proportion of alternate reads. Under the null hypothesis that the reads are generated due to amplification artefact, *p =* 0.125. Under the alternate hypothesis that the reads are generated due to a true mosaic variant, *p =* 0.125 +*θ* where *p* is the allelic balance estimated by kernel smoothing^24^. Since the models of the two hypotheses are nested, −2 × (log/(*X;* 0.125 +*θ)/f(X;* 0.125)) ~ *X*_1_^2^. We applied this model at all 400 sites at which we attempted amplicon validation across the 12 deeply sequenced single cells to obtain high quality single cell genotypes. Scripts for running this model are available at https://github.com/parklab/SCGenotypermini.

#### Variant validation using 10X linked-read data (*contributed by Yifan Wang and Jeffrey Kidd*)

We used haplotype information provided by 10X Genomics linked-read sequencing data to eliminate false positive mosaic SNV calls and to provide support for *bona fide* mosaic SNVs. Briefly, the 10X Genomics linked-read approach is a barcoded short-read sequencing technology, where individual hMw genomic DNA molecules (~50 kb) initially are attached to a bead then incorporated into droplets. Within each droplet, the HMW DNA fragments are fragmented into small pieces, assigned a unique barcode, and then undergo amplification. The resultant DNA libraries from each individual droplet then were subject to standard Illumina short-read sequencing. Short reads containing the same barcodes were aligned and re-assembled into larger linked-reads using the LongRanger pipeline (https://support.10xgenomics.com/genome-exome/software/pipelines/latest/what-is-long-ranger). Overlapping individual HMW fragments that shared informative germline SNP alleles were used reconstruct longer, continuous haplotype segments. This results in the assignment of sequencing reads derived from the initial HMW fragments to individual haplotypes. Using this information, we identified and removed candidate mosaic SNVs that were found on both parental haplotypes, which likely arose from systematic sequencing errors. Moreover, the haplotype information could be used to identify candidate mosaic SNVs that were represented >90% of the reads on a single parental haplotype and are likely germline SNPs. By comparison, *bona fide* somatic SNVs are detected as a minor allele on a single parental haplotype. Tools for analyzing linked-read BAMs are available at: https://github.com/KiddLab/hapfilter-10X

#### Variant validation using ddPCR (*contributed by Reenal Pattni*)

We performed ddPCR using the primers and probe listed in **Table S6**. We initially designed 29 ddPCR assays to assess candidate mosaic SNVs present at <3% VAFs and/or near know SVs (**Table S3**).This initial set included: (i) 4 mosaic SNVs at higher VAFs (12%-28% in amplicon sequencing); (ii) 5 mosaic variants at lower VAFs (1%-2.5% in amplicon sequencing); (iii) 3 SNVs deemed to be germline SNVs; and (iv) 17 false-positive calls (including 6 that were present at <0.5% VAFs). The ddPCR reaction mixture (20 μL) contained 2 μL of bulk gDNA or NA12878 (25ng) template, 250 nM of each primer (reference and assay specific pairs), and 250 nM TaqMan probe (Ref-HEX and assay specific-FAM dye labeled) in a 1× Bio-Rad Supermix (Bio-Rad, Hercules, CA, USA). The 20μl PCR reaction was mixed with Bio-Rad droplet generator oil and partitioned into 15,000–20,000 droplets using the Bio-Rad QX-100 droplet generator (Bio-Rad). The generated droplets were transferred to a 96-well PCR reaction plate and sealed with a pierceable sealing foil. PCR conditions were 10 min at 95°C, 40 cycles of denaturation for 30 s at 94°C, assay specific anneal/extension temperatures 50°C-57.5°C (as noted in table) for 60 s with ramp rate of 2.5°C s^−1^, followed by 10 min at 98°C and a hold at 4°C. Post-PCR amplification each droplet was checked for fluorescence to count the number of droplets that yielded positive/negative results, using the Bio-Rad QX-100 droplet reader (Bio-Rad) and frequencies calculated based on total droplet counts per well. In the end, we were able to design informative ddPCR validation tests for 13 candidate mosaic SNVs. For the remaining 16 variants, we could not design effective probes because the variants were within AT rich or repetitive regions or the assay failed to separate clusters at various annealing temperatures.

#### Validation result consolidation (*contributed by Yifan Wang and Alexej Abyzov*)

We tested 383 sites using PCR amplicon-based sequencing (Fig. 3). Of these, 280 were determined to be false positive calls, 64 had read support above background levels (eventually called as 9 germline SNVs, 17 false positive calls, and 38 true mosaic SNVs), and 39 had a discordant validation status between two different BSMN groups (eventually called as 2 germline SNVs, 32 false positive calls, and 5 true mosaic SNVs).

To further evaluate the discordant calls, we considered experiments yielding overlapping end sequence reads as more reliable because they resulted in higher quality sequencing data (*i.e.*, higher confidence base calls, reduced indels, and better mappability); these analyses allowed us to resolve 11/39 candidates. Other information that allowed the interrogation of the remaining calls included: (1) determining the number of haplotypes observed in 10X genomics linked-read sequencing datasets (*i.e.*, “3 haplotypes” were observed for true positive calls and “4 haplotypes” were observed for false positive calls); (2) eliminating SNVs at a <0.005 VAF in the PCR amplicon-based sequencing experiments (as such variants are unlikely be called from WGS data); (3) eliminating calls with fewer than 3 reads per WGS replicate; (4) eliminating variants within homopolymer tracts, CNVs, within 3 bps of indels, and near a germline Alu insertion (**Fig. S8**). We also applied the above criteria to invalidate 17/64 candidates with read support above background levels. Furthermore, 9/64 and 2/39 discordant candidates were germline SNVs, as their VAF approached 0.5 and only “2 haplotypes” were observed for the candidate calls in 10X linked-read sequencing data. Following ddPCR, one SNV candidate initially assigned as a false positive was reassigned as a true SNV. Finally, based on the examination of 10X linked-read sequencing datasets, one of the 17 candidates that lacked enough data from the PCR amplicon-based sequencing experiments (“NED”; not enough data) was determined to be a germline SNV. By consolidating all validation results, we arrived at a confident set of 43 mosaic SNVs.

#### Panel of normal (PON) mask using 1000 Genomes 2504 whole genome sequencing libraries (*contributed by Yifan Wang and Jeffrey Kidd*)

We applied a PON mask for all the candidate somatic SNV sites to exclude the possible sequencing artifacts using the 1000 Genomes Project whole genome sequencing libraries. We collected the counts for reference alleles and alternative alleles for each candidate site in each of the 2504 samples with a read quality cutoff at 20 and a base quality cutoff at 20. Candidate sites that have greater than 0.05 candidate allele frequency in more than 5 samples were identified as false positives. Thus, the PON mask filter out possible recurrent artifacts that are induced during sequencing or library preparation.

#### Classifying false-positive calls (*contributed by Max Sherman*)

We consolidated the above filtering strategies into a naïve Bayes classifier and assessed our ability to characterize the 400 putative candidate mosaic SNVs using a “leave-one-out” cross-validation method. Briefly, we iteratively trained the model on validation results from 399 mosaic SNVs and predicted the validation status of the held-out SNV. This classifier uses features such as the allele count of a candidate somatic SNV across the 2504 samples in the 1000 Genomes Project as well as Chromium 10X linked read and single cell properties to aid in the discrimination of true somatic SNVs from false-positive calls (see below). Given that trade-offs between maximizing the number of true-positive calls and excluding false-negative calls are inevitable in statistical modeling, we ultimately arrived at conservative calling parameters, where the classifier achieved 90% precision, 42% sensitivity, and 99% specificity. While this model suffers with respect to sensitivity, the chosen values maximize the rejection of false-positive calls at the expense of removing some false-negative calls (**Table S4**). For example, false-negative calls were mostly present at low VAFs (median = 0.013) and lacked haplotype support in Chromium 10X linked-read (*i.e.*, 15/25 false-negative calls lacked haplotype support, whereas only 4/18 true-positive calls lacked haplotype support) and/or single cell sequencing datasets (*i.e.*, 22/25 falsenegative calls lacked support in single cell datasets as compared to 9/18 true-positive calls). Overall, these results were consistent with our prior observations and indicated that integrating various sequence characteristics can distinguish between *bona fide* mosaic SNVs and false-positive signals. Indeed, these criteria, as well as additional filtering strategies, have been consolidated for use in a recently developed machine-learning algorithm, MosaicForecast^32^.

#### Naive Bayes classifier combining validation approaches (*contributed by Max Sherman*)

We trained a naïve Bayes classifier to provide the probability that a given site will pass or fail amplicon validation based on the properties of the site in the 75 WGS panel of normal samples, 10X sequencing from the common brain tissue, and single cell whole-genome sequencing. Given a set of training sites, we calculated the conditional likelihood of a site having the following properties given its validation status (pass or fail):

- Panel of normal properties:

○ Presence of alternate reads in ≤ 5 of the 75 panel of normal samples
○ Presence of alternate reads in >5 of the 75 panel of normal samples
- 10X properties:

○ PASS 10X filters (see above Methods)
○ FAIL 10X filters for any reason
○ N10X information available due tlack of alternate reads
- Single cell properties:

○ Nalternate reads present in any single cells
○ Alternate reads present in <4 single cells
○ Alternate reads present in ≥4 single cells
○ Locus flagged as PASS by single cell genotyping method
○ Locus flagged as LOWQUAL by single cell genotyping method
○ Locus flagged as ARTIF by single cell genotyping method
○ Locus has nflag assigned by single cell genotyping algorithm

The conditional likelihood was calculated as the number of sites exhibiting that property and having a certain validation status divided by the number of sites with that validation status. To prevent events of probability zero, each category was given a prior pseudocount of 1. The prior probability of passing (failing) was calculated as the number of test sites in the training data which passed (failed) amplicon validation divided by the number of test sites in the training data.

Let *B* be a binary random variable which takes on the value 0 if the site fails validation and 1 if it passes validation. Let S be the set of properties given above for some new locus:

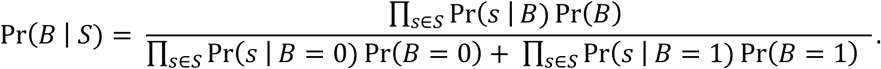

The sensitivity and specificity of this approach was calculated using leave-one-out cross-validation on the 400 sites.

#### Implementation of the best practices workflow (*contributed by Taejeong Bae*)

The best practices workflow was implemented as a computational pipeline composed of a series of bash job scripts compatible with the job scheduler and Sun Grid Engine. The software includes the steps of uniform data processing, variant calling with GATK at various ploidies, and variant filtering. It is accessible at: https://github.com/bsmn/bsmn-pipeline. Although the pipeline is executable at any cluster system using Sun Grid Engine as a job scheduler, we used AWS ParallelCluster (https://github.com/aws/aws-parallelcluster) as the running environment for the BSMN project. The configuration used for setting up the ParallelCluster is available at: https://github.com/bintriz/bsmn-aws-setup.

#### Cell lineage tree reconstruction (*contributed by Taejeong Bae*)

To reconstruct a cell lineage tree from single cell data, we examined whether the union set of 49 SNVs identified in our validation call set and by the application of the best practices workflow to DLPFC replicate 6 WGS were shared among twelve NeuN+ neuron single cell datasets. An SNV was considered “present” if we could find at least one supporting read in a single cell. The VAFs of the 49 SNVs also were estimated from the following WGS datasets: DLPFC (4 samples), NeuN+ (2 datasets) and NeuN-(1 dataset) DLPFC cell fractions, cerebellum (1 dataset), dura mater (1 dataset), and dural fibroblasts (2 datasets). We conducted hierarchical clustering of SNVs based upon their VAFs in different tissues using average linkage with Canberra distance. Clustering of single cells was performed by assessing shared genotypes using Jaccard distance. The final lineage tree was constructed by manual inspection of shared genotypes in each cell and the SNV VAF values across tissues. SNVs defining earlier developmental lineages have higher VAFs than SNVs defining later developmental sub-lineages. Six SNVs with highest VAFs clustered together and were present in two distinct groups of single cells (the L1 and L2 lineages). We found that 4/6 SNVs (SNVs 1-4) denote the L1 lineage and 2/6 SNVs (SNVs 10 and 11) denote the L2 lineage. This analysis was conducted using the SciPy package of python. Notably, one single cell dataset lacked the 49 SNVs and was excluded from the analysis.

Two calls (SNV6 and SNV16) were not validated in our PCR amplicon-based sequencing experiments, but passed the 10X Genomics linked-read haplotype test. Further analyses indicated that SNV6 and SNV16 exhibited consistent VAFs across multiple sequencing replicates, that no apparent alignment or sequencing artifact artifacts surround the SNV site, and that these SNVs defined sub-branches of L1 and L2 lineages. Moreover, SNV16 appeared together with SNV17 in a single cell and had similar VAFs, suggesting they represent markers for a sub-branch of the L2 lineage and, consequently, SNV16 is a bona fide mosaic SNV.

#### Predicting early cell lineages from bulk DNA (contributed by Taejeong Bae)

To identify mosaic SNV pairs that putatively were marking different cell lineages in WGS bulk DNA, we calculated all possible pairwise SNV VAF anti-correlation values for the 49 somatic mosaic SNVs identified in our validation set and best practices pipeline. For each pairwise comparison, we calculated the mean distance of VAFs of the SNV pair across multiple tissue samples according to the diagonal line: VAF1 + VAF2 = 0.5. We then used that distance as a score to rank SNV pairs by how well they fit the diagonal line (the lower score is better). The eight best scores corresponded to sNv pairs, where one represented the L1 lineage and the other one represented the L2 lineage. For seven pairs the scores were differed significantly (<0.05) from the other tested pairs (**Fig. S11**).

