## supplementary material for "Comprehensive identification of somatic nucleotide variants in human brain tissue"

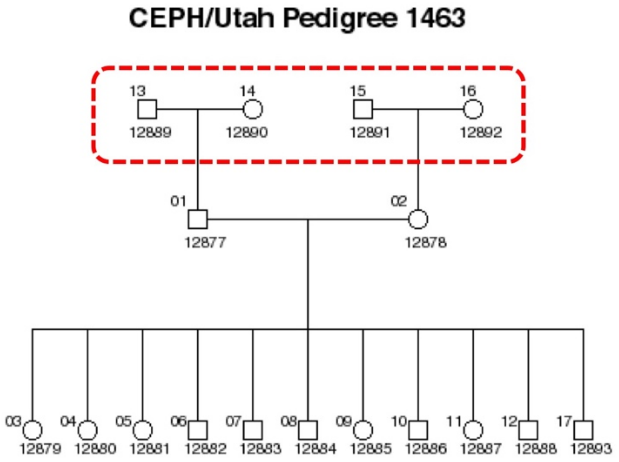


| Genome | Mix1 | Mix2 | Mix3 | *Mix1 + Mix2* |
|---|---|---|---|---|
| NA12889 | 4% | 2% | 0% | 3% |
| NA12891 | 8% | 4% | 0% | 6% |
| NA12890 | 16% | 8% | 0% | 12% |
| NA12892 | 72% | 86% | 100% | 79% |
| **Depth** | **~130X** | **~110X** | **~100X** | **~240X** |

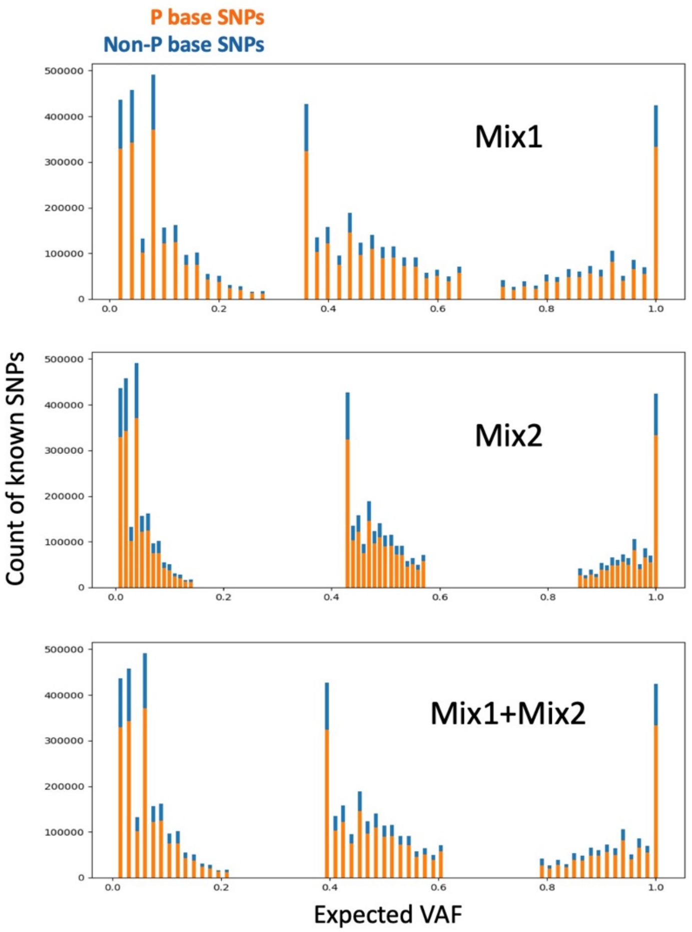


**Supplementary Figure 1: Additional information about the samples used in DNA mixing experiments to simulate mosaic SNVs.** Left: The genetic pedigree of four individuals (red dashed lines) and DNA dilutions used in the mixing experiments. Right: Histograms depicting the expected VAFs of simulated mosaic SNVs in the different mixes.

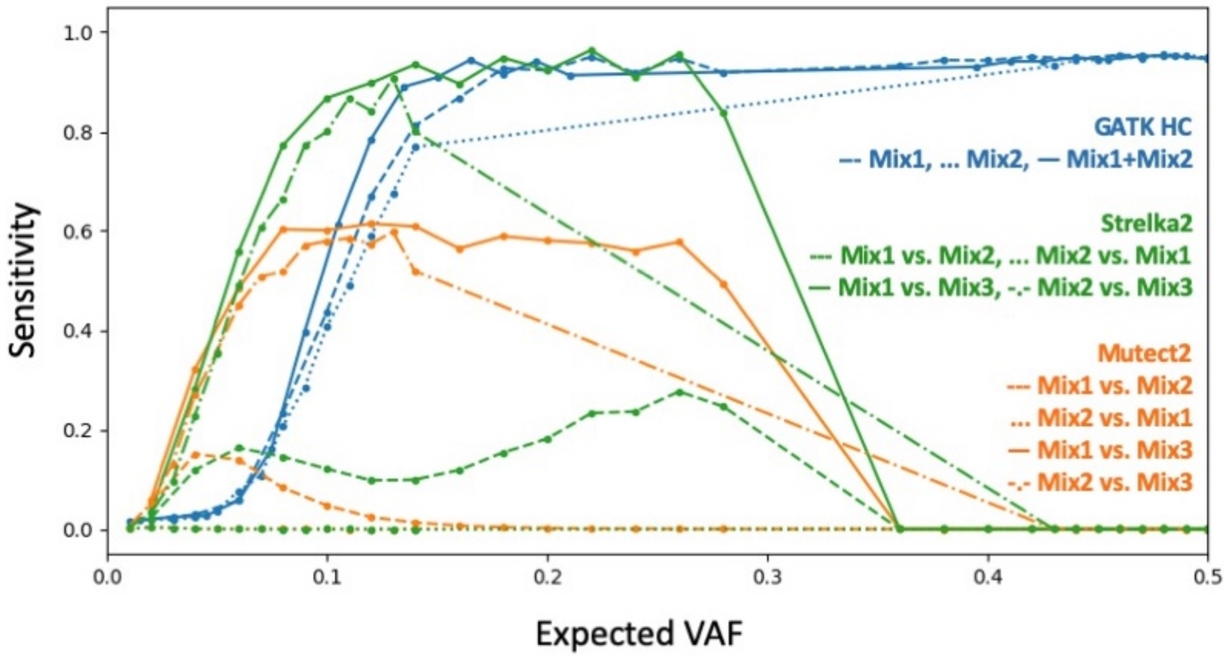


**Supplementary Figure 2: The use of existing tools to detect simulated mosaic SNVs.** The expected VAFs of simulated mosaic SNV (x-axis) and the sensitivity of existing tools (y-axis) to detect simulated SNVs in the DNA mixing experiments. The GATK Haplotype Caller was applied in single sample mode; Strelka2 and MuTect2 were applied in paired sample mode.

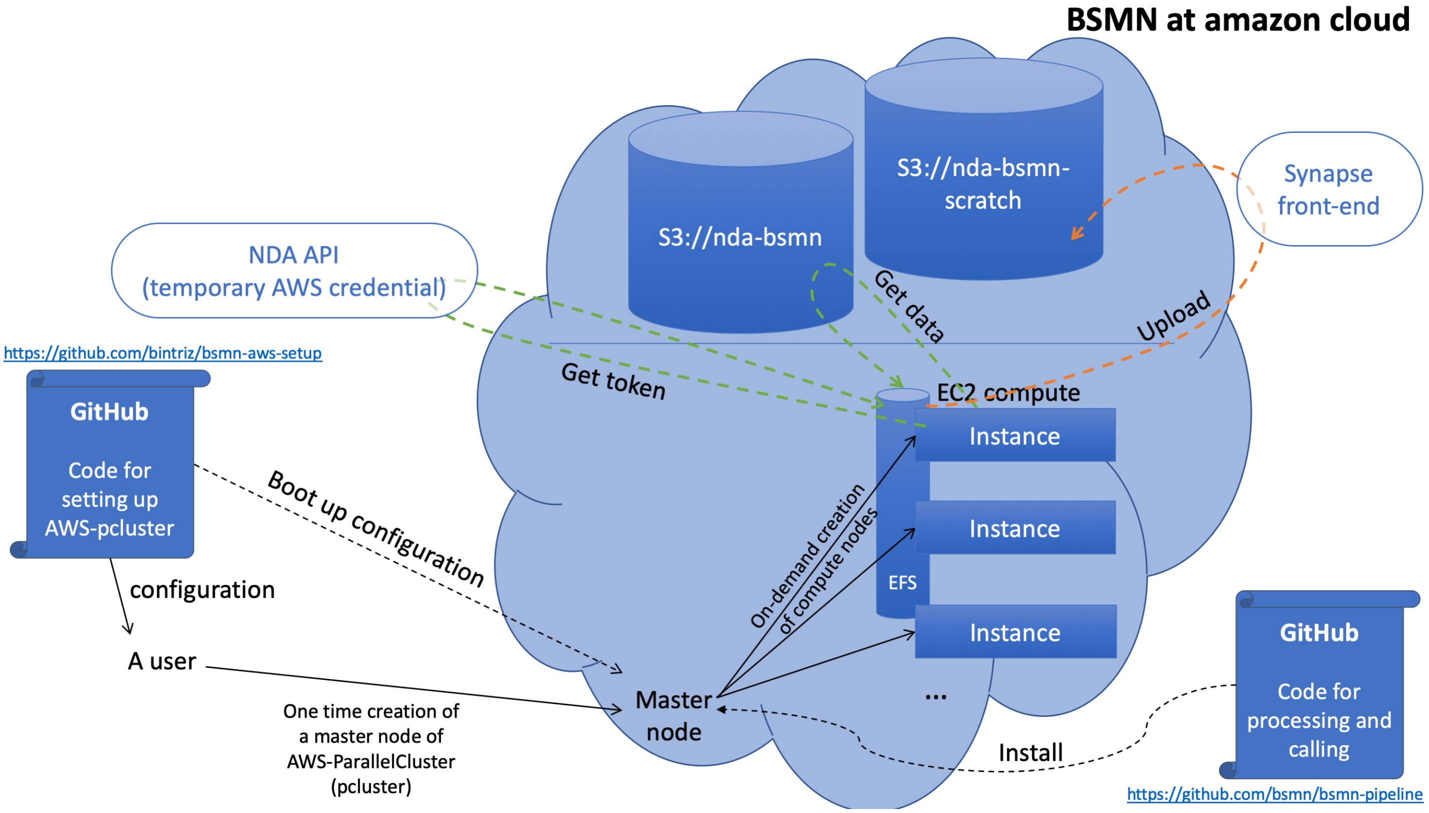


**Supplementary Figure 3: A schematic depicting the common processing workflow using the Amazon Web Service (AWS) cloud computing platform.** The generated raw data files (.bam or .fastq formats) from each BSMN node were uploaded onto the shared BSMN S3 bucket. The AWS ParallelCluster was used to create a Sun Grid Engine (SGE) job queuing system. The cluster also was configured to have an attached Elastic File System (EFS) to function as a shared file system. We then installed the BSMN common pipeline (https://github.com/bsmn/bsmn-pipeline). Upon downloading each sample, the pipeline assigns a temporary S3 bucket access token based upon the users NDA credential, which enables access to the data collection. At the end of processing, the resultant .cram file for each sample was uploaded to the BSMN shared scratch S3 bucket using Synapse.

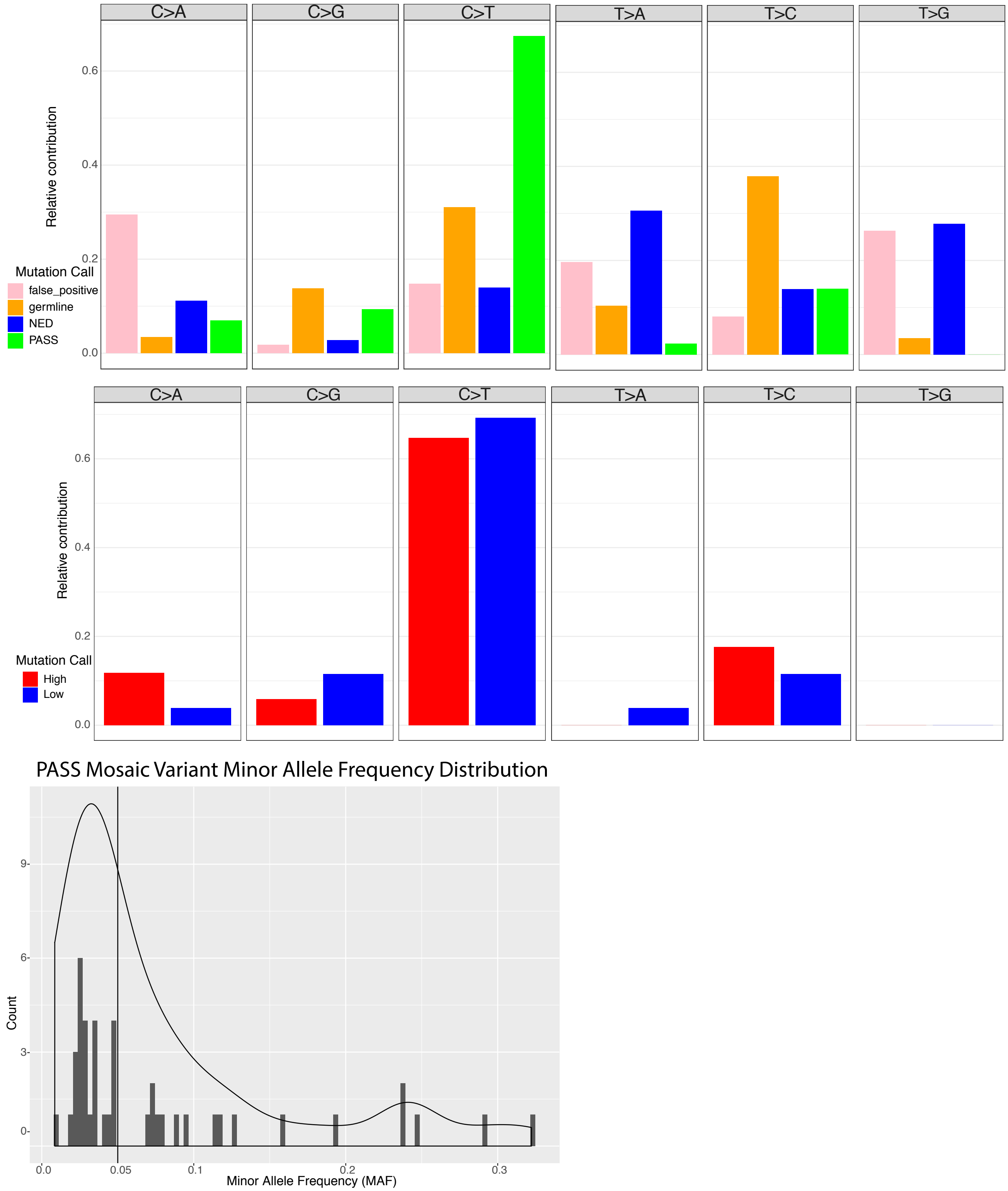


**Supplementary Figure 4: Mutation spectra for the 400 candidate mosaic SNV calls subject to validation analyses.** Top: Mutation spectra for true-positive, false-positive, germline, and ambiguous calls. Middle: Mutation spectra for validated mosaic SNVs with the highest (blue bars; 21 SNVs) and lowest (red bars; 22 SNVs) VAFs. There is no significant difference between the mutation spectra for the high and low VAF calls. Bottom: VAF distributions for validated mosaic SNVs in the common reference brain.

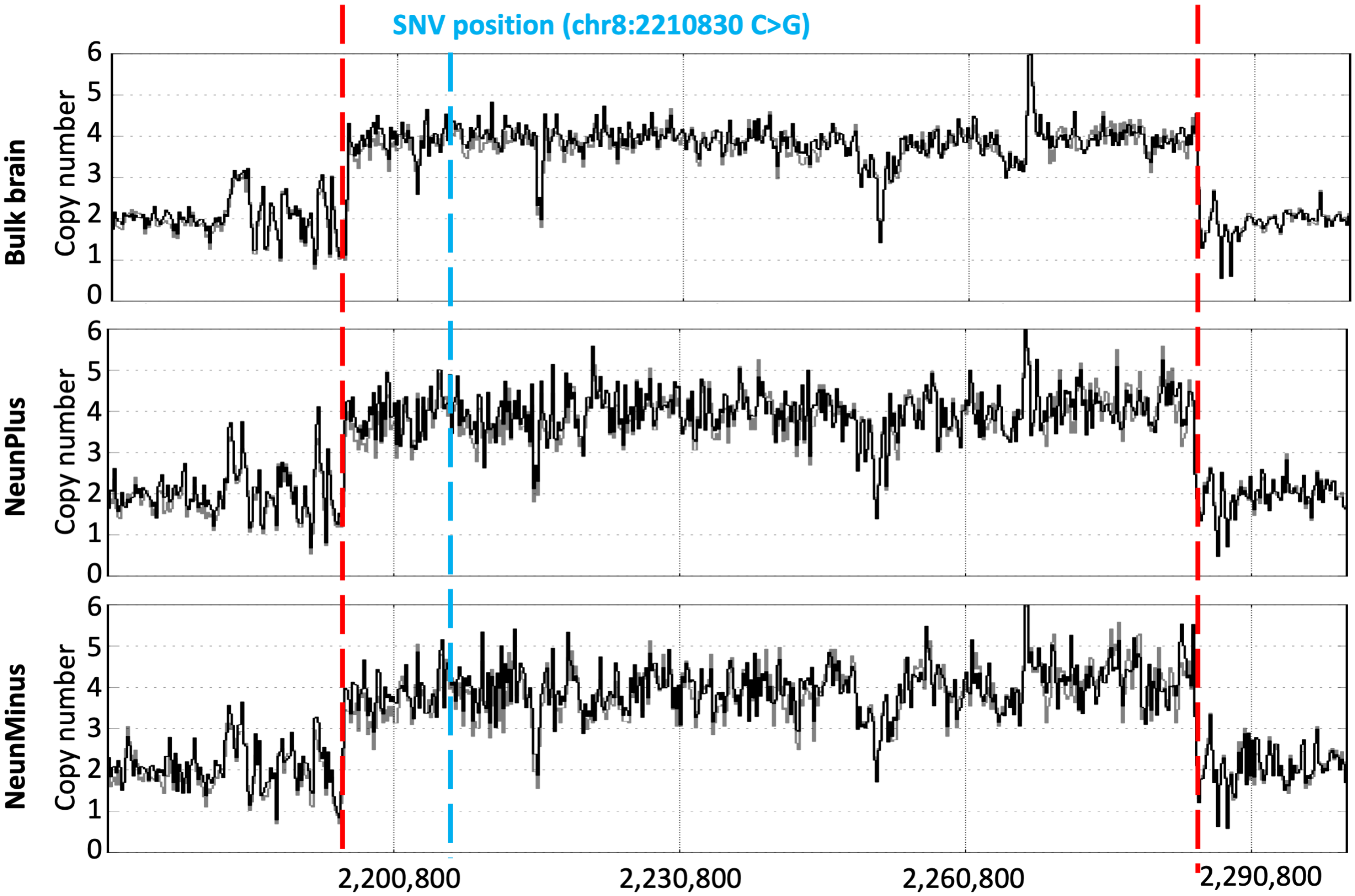


**Supplementary Figure 5: An illustrative example of a false-positive mosaic SNV call located within a genomic region containing a copy number gain in three different brain samples (*i.e.*, bulk brain DNA, NeuN+, and NeuN- cells).** The black line indicates sequencing read depth. The read vertical dashed lines denote the germline copy number gain boundaries. The blue vertical dash line shows the position of the candidate SNV. In this example, a germline SNP (marked with blue vertical dash line) within the CNV region led to the false-positive call.

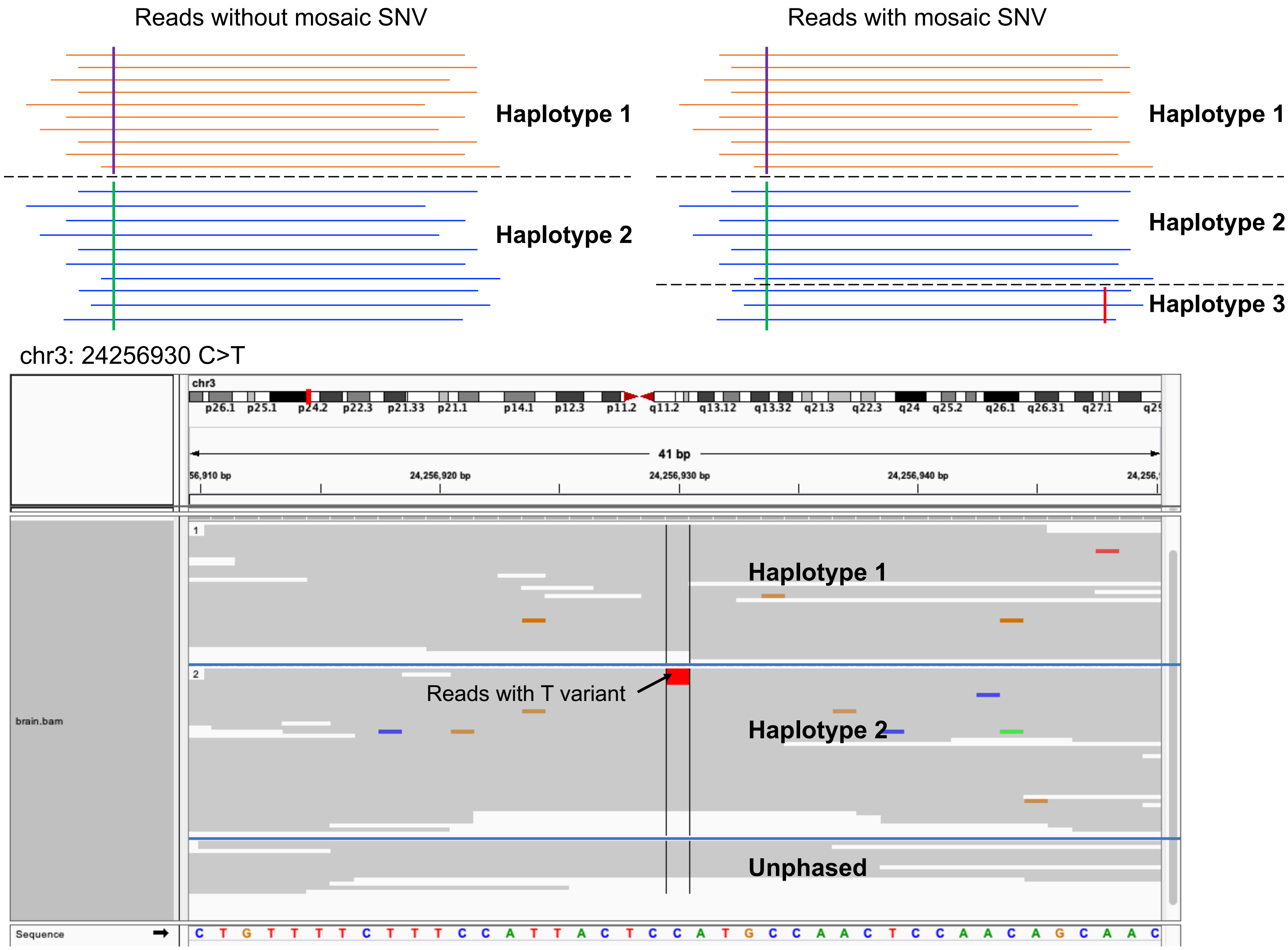


**Supplementary Figure 6: An example of true moisac SNV supported by Chromium 10X linked-read data.** Top left: Schematic representation the two parental haplotypes (orange and blue horizontal lines, respectively) containing a distinguishing germline SNP. The purple vertical line depicts the germline SNP in haplotype 1; the green vertical line depicts germline SNP in haplotype 2. Top right: *bona fide* mosaic SNV calls result in three haplotypes. Orange horizontal lines represent haplotype 1; blue horizontal lines represent haplotype 2; blue horizontal lines with red vertical bar depict the alternative mosaic SNV allele (haplotype 3), which only are present on a subset of haplotype 2 reads. Bottom: Read level data from the Chromium 10X dataset that allow the identification of a *bona fide* somatic mosaic SNV. Reads in haplotype 1 only have the reference C allele, whereas a fraction of reads in haplotype 2 have an alternative T allele. The unphased reads could not be assigned to specific haplotypes.

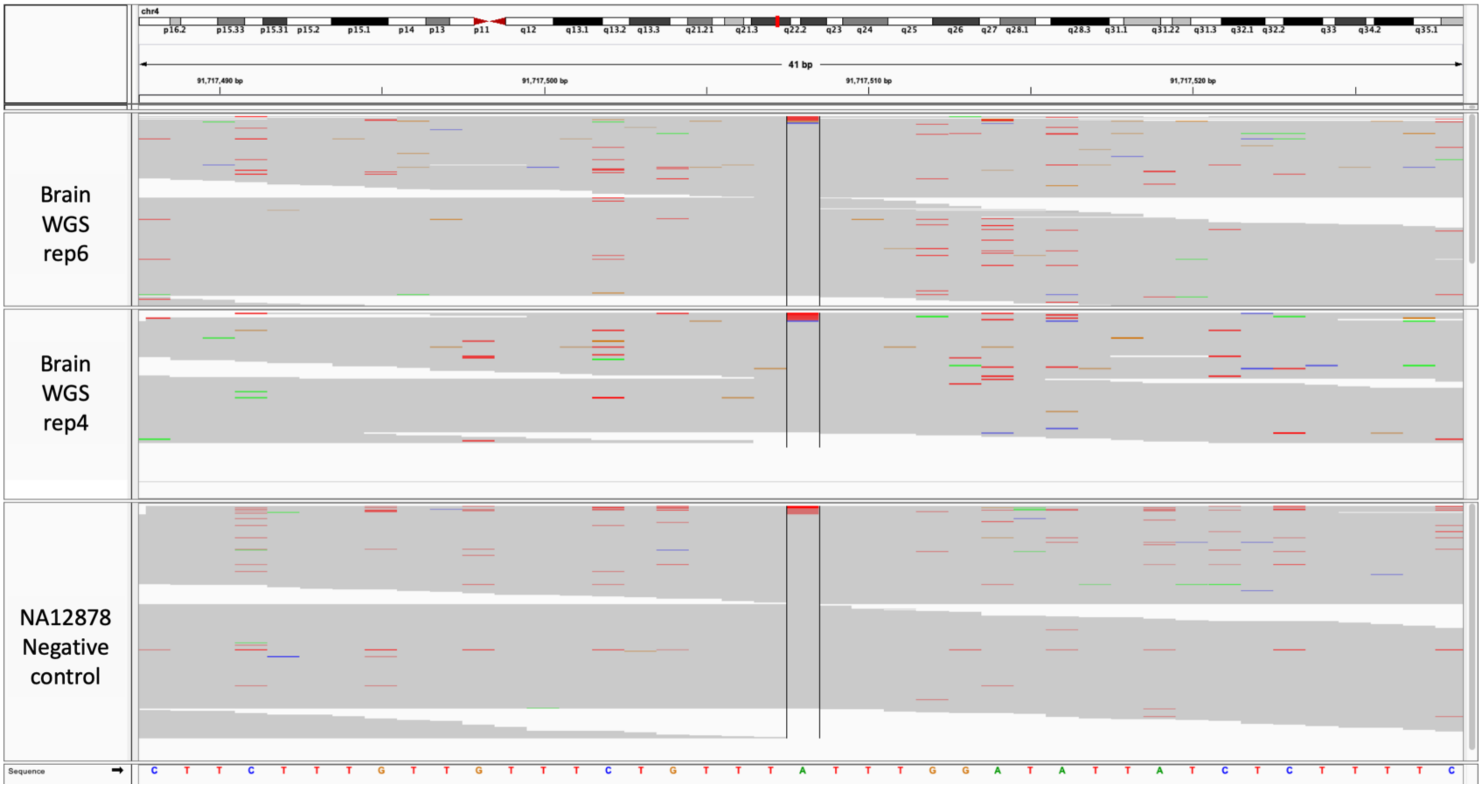


**Supplementary Figure 7: An illustrative example of a false-positive mosaic SNV located within a short homopolymeric repeat.** A putative A>T mosaic SNV call was made in two replicates for an A nucleotide (shown in green) that resides in six T nucleotides (shown in red). Evidence for the same putative mosaic SNV call also was observed in NA12878 control DNA. This recurrence suggests that the A to T difference represents a false-positive call, which likely represents a systematic sequencing error.

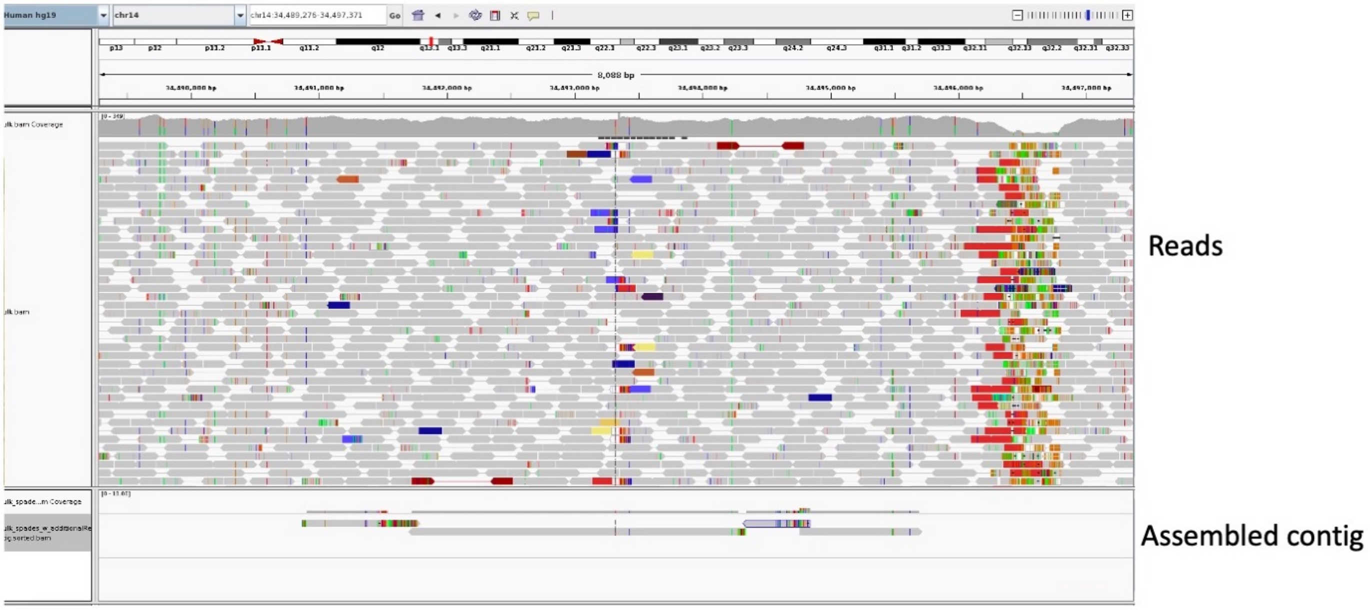


| ACTIONS | QUERY | SCORE | START | END | QSIZE | IDENTITY | CHROM | STRAND | START | END | SPAN |
|---|---|---|---|---|---|---|---|---|---|---|---|
| browser details | chr14_event | 2548 | 1 | 2869 | 2926 | 99.2% | chr14 | + | 34491725 | 34494279 | 2555 |
| browser details | chr14_event | 322 | 1598 | 1942 | 2926 | 98.3% | chrX | + | 70080722 | 70081223 | 502 |
| browser details | chr14_event | 310 | 1625 | 1989 | 2926 | 99.1% | chr1 | + | 231303065 | 231303694 | 630 |

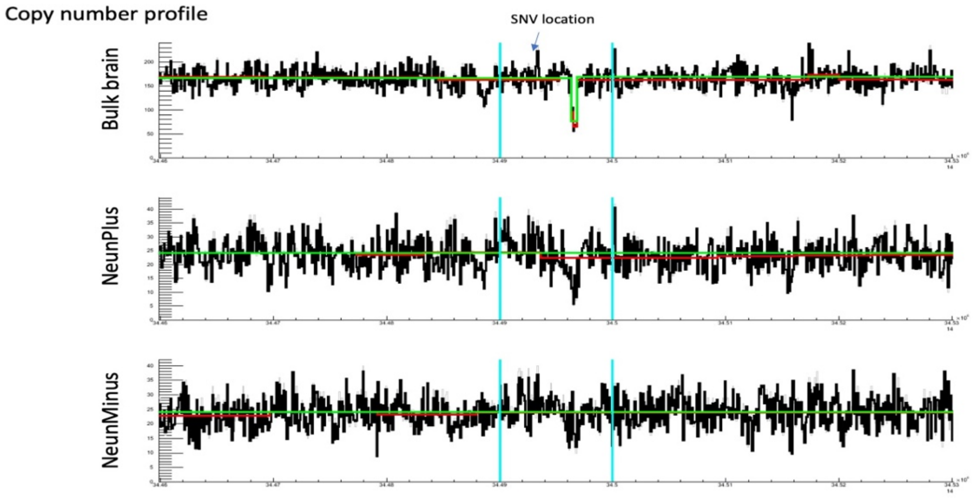


**Supplementary Figure 8: An illustrative example of a false-positive somatic SNV call near a genomic structural variant (*i.e.*, a polymorphic Alu insertion).** Top: Multiple read pairs (blue, yellow, and red rectangles) had one an end that aligned to the displayed locus (gray dashed vertical line) on chromosome 14 and another end that aligned to a different chromosome. Middle: The assembled read contig contained a ~300 bp full-length human-specifc Alu insertion (from the Ya5 or Yb8 subfamily) that was not present in the UCSC genome browser. Secondary alignments to the X chromosome and chromosome 1 revealed evidence for the presence of a full-length Alu element (second and third browser track lines). Bottom: The region harboring the false-positive mosaic SNV call does not reside within the polymorphic Alu element.

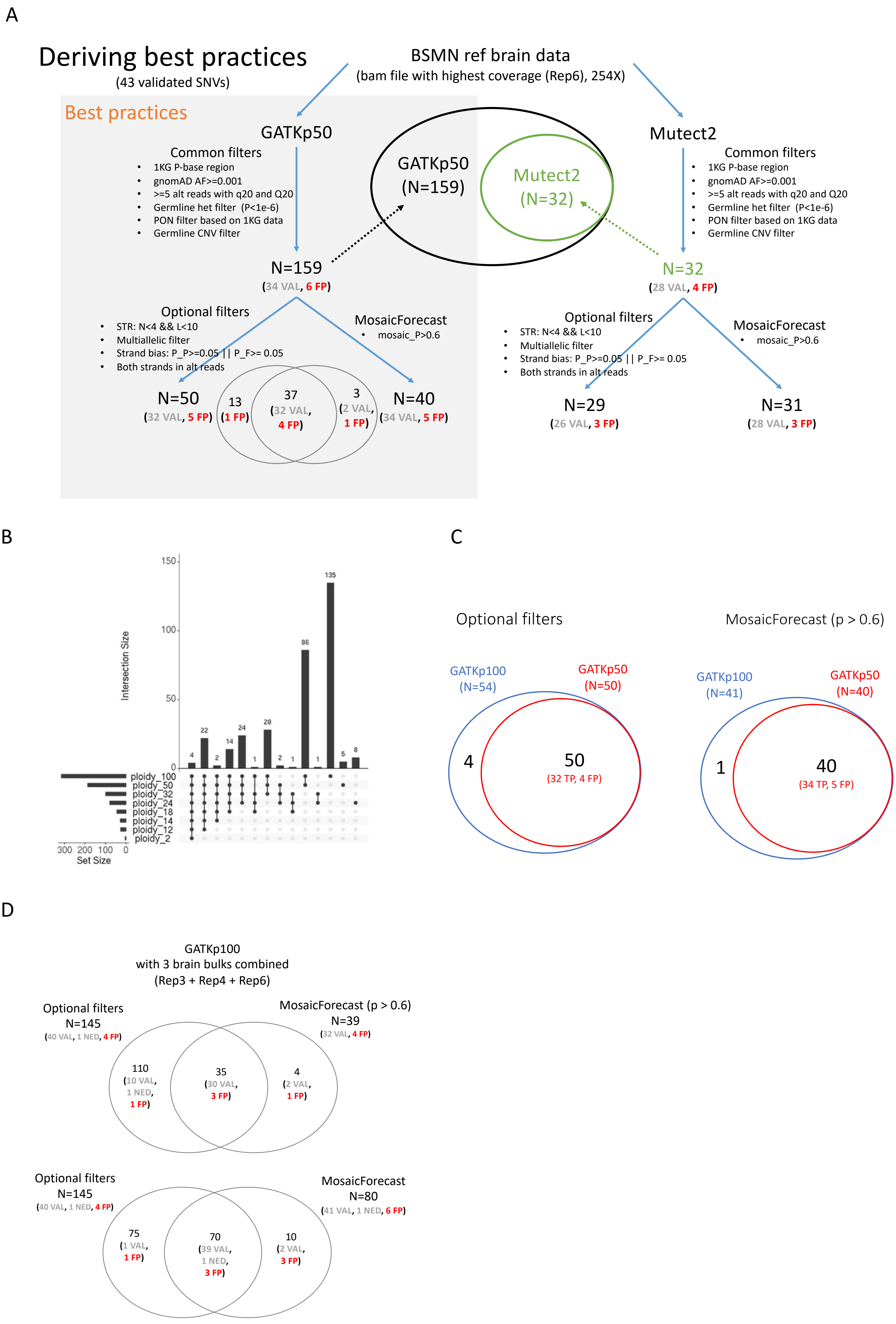
A
Deriving best practices
(43 validated SNVs)
BSMN ref brain data
(bam file with highest coverage (Rep6), 254X)
Best practices
GATKp50
Common filters
1KG P-base region
gnomAD AF>=0.001
>=5 alt reads with q20 and Q20
Germline het filter (P<1e-6)
PON filter based on 1KG data
Germline CNV filter
N=159
(34 VAL, 6 FP)
GATKp50
(N=159)
Mutect2
(N=32)
Mutect2
Common filters
1KG P-base region
gnomAD AF>=0.001
>=5 alt reads with q20 and Q20
Germline het filter (P<1e-6)
PON filter based on 1KG data
Germline CNV filter
N=32
(28 VAL, 4 FP)
Optional filters
STR: N<4 && L<10
Multiallelic filter
Strand bias: P_P>=0.05 || P_F>= 0.05
Both strands in alt reads
MosaicForecast
mosaic_P>0.6
N=50
(32 VAL, 5 FP)
13
(1 FP)
37
(32 VAL,
4 FP)
3
(2 VAL,
1 FP)
N=40
(34 VAL, 5 FP)
Optional filters
STR: N<4 && L<10
Multiallelic filter
Strand bias: P_P>=0.05 || P_F>= 0.05
Both strands in alt reads
MosaicForecast
mosaic_P>0.6
N=29
(26 VAL, 3 FP)
N=31
(28 VAL, 3 FP)
B
Intersection Size
150
100
50
0
4
22
2
14
24
1
28
2
1
86
1
135
5
8
ploidy_100
ploidy_50
ploidy_32
ploidy_24
ploidy_18
ploidy_14
ploidy_12
ploidy_2
300 200 100 0
Set Size
C
Optional filters
GATKp100
(N=54)
GATKp50
(N=50)
4
50
(32 TP, 4 FP)
MosaicForecast (p > 0.6)
GATKp100
(N=41)
GATKp50
(N=40)
1
40
(34 TP, 5 FP)
D
GATKp100
with 3 brain bulks combined
(Rep3 + Rep4 + Rep6)
Optional filters
N=145
(40 VAL, 1 NED, 4 FP)
MosaicForecast (p > 0.6)
N=39
(32 VAL, 4 FP)
110
(10 VAL,
1 NED,
1 FP)
35
(30 VAL,
3 FP)
4
(2 VAL,
1 FP)
Optional filters
N=145
(40 VAL, 1 NED, 4 FP)
MosaicForecast
N=80
(41 VAL, 1 NED, 6 FP)
75
(1 VAL,
1 FP)
70
(39 VAL,
1 NED,
3 FP)
10
(2 VAL,
3 FP)

E

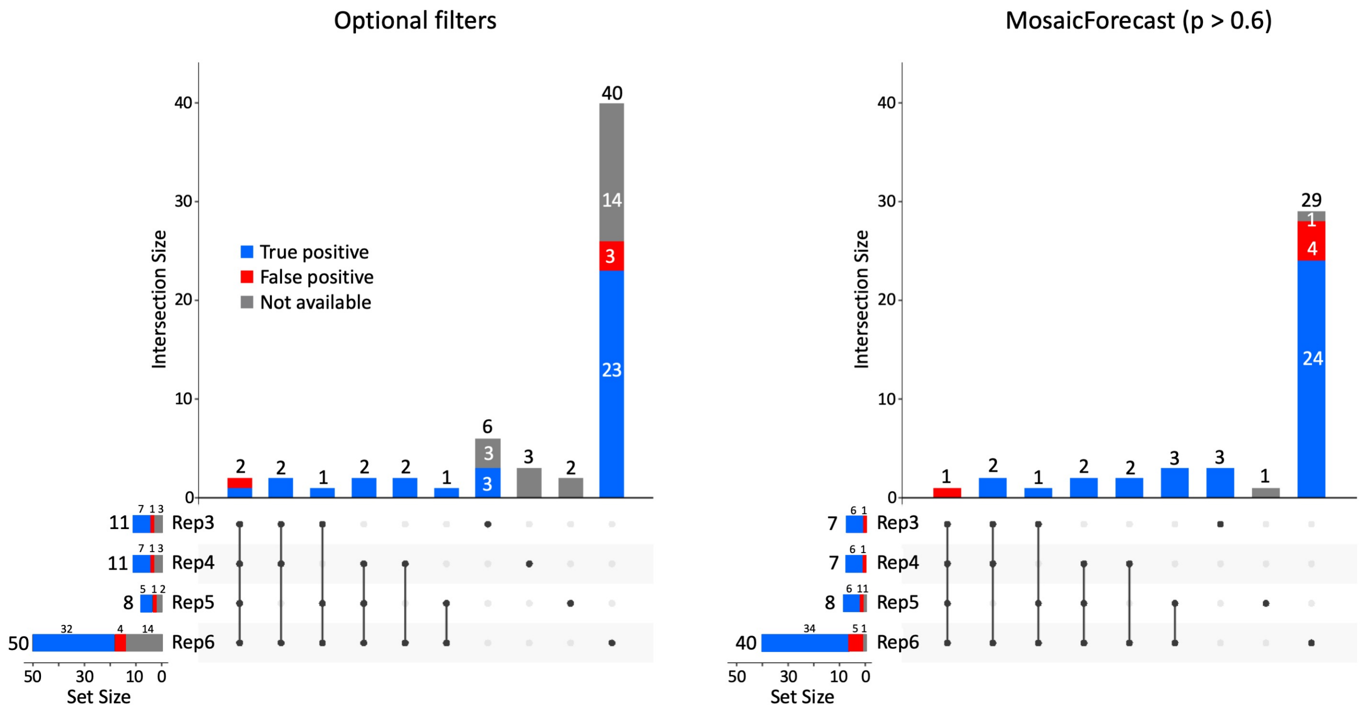


**Supplementary Figure 9: The application of the "best practices" workflow for calling candidate mosaic SNVs.** *(A)* The GATK haplotype caller (using a ploidy setting of 50) and MuTect2 were used to call candidate mosaic SNVs in a 250X WGS dataset**.** The use of GATK with a ploidy setting of 50 leads to more sensitive candidate SNV discovery with similar specificity when compared to MuTect2. The application of MosaicForecast consistently results in a higher percentage of validated calls when compared to analyses conducted using the "optional filters" strategy. NED stands for "not enough data" (see text). MosaicForecast was applied with score cutoff of 0.6 (i.e., p > 0.6). *(B)* The detection of candidate mosaic SNVs using different GATK ploidy settings. A GATK ploidy setting of 100 only yielded a marginal increase in mosaic SNV calls when compared to a ploidy setting of 50. *(C)* Venn diagrams comparing the results of SNV variant filtering strategies using either the "optional filters" or MosaicForecast setting using GATK ploidy settings of 50 and 100, respectively. Using the ploidy setting of 100 only leads to a marginal increase in the number of candidate mosaic SNV calls. *(D)* Increasing WGS sequencing coverage by combining multiple replicates (replicate 5 was excluded because of poor sequence quality) results in a greater number of candidate somatic mosaic SNV calls. *(E)* UpSet plots depicting the numbers of somatic SNVs detected using the "best practices" workflow using the "optional filters" (left) and the MosaicForecast (right) pipelines. The calls were generated from the four listed WGS replicates. The percentage of validated calls is higher when relying on MosaicForecast.

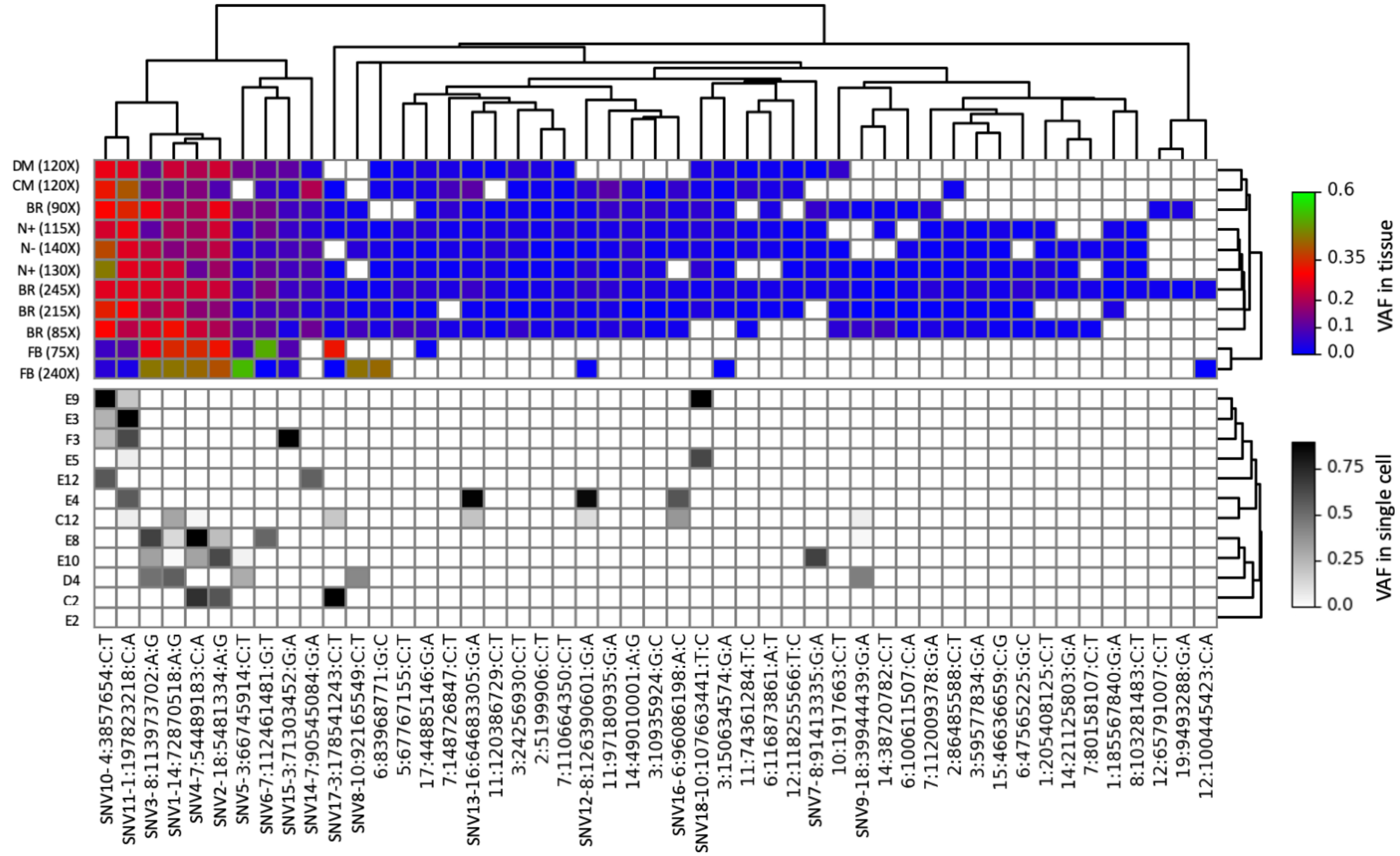


**Supplementary Figure 10: Validated somatic SNV VAFs in bulk WGS and single cell sequencing datasets.** Top: Datasets and fold sequencing coverage are indicated on the left. The VAF color key (multi-colored rectangle) is indicated on the right. Many of the validated somatic SNVs are present at low VAFs. Bottom: Single-cells are indicated on the left. The VAF color key (gray rectangle) is indicated on the right. Due to uneven amplification of single cell genomes, support for a candidate mosaic SNV variant can vary from significantly both below and above a 0.5 VAF.

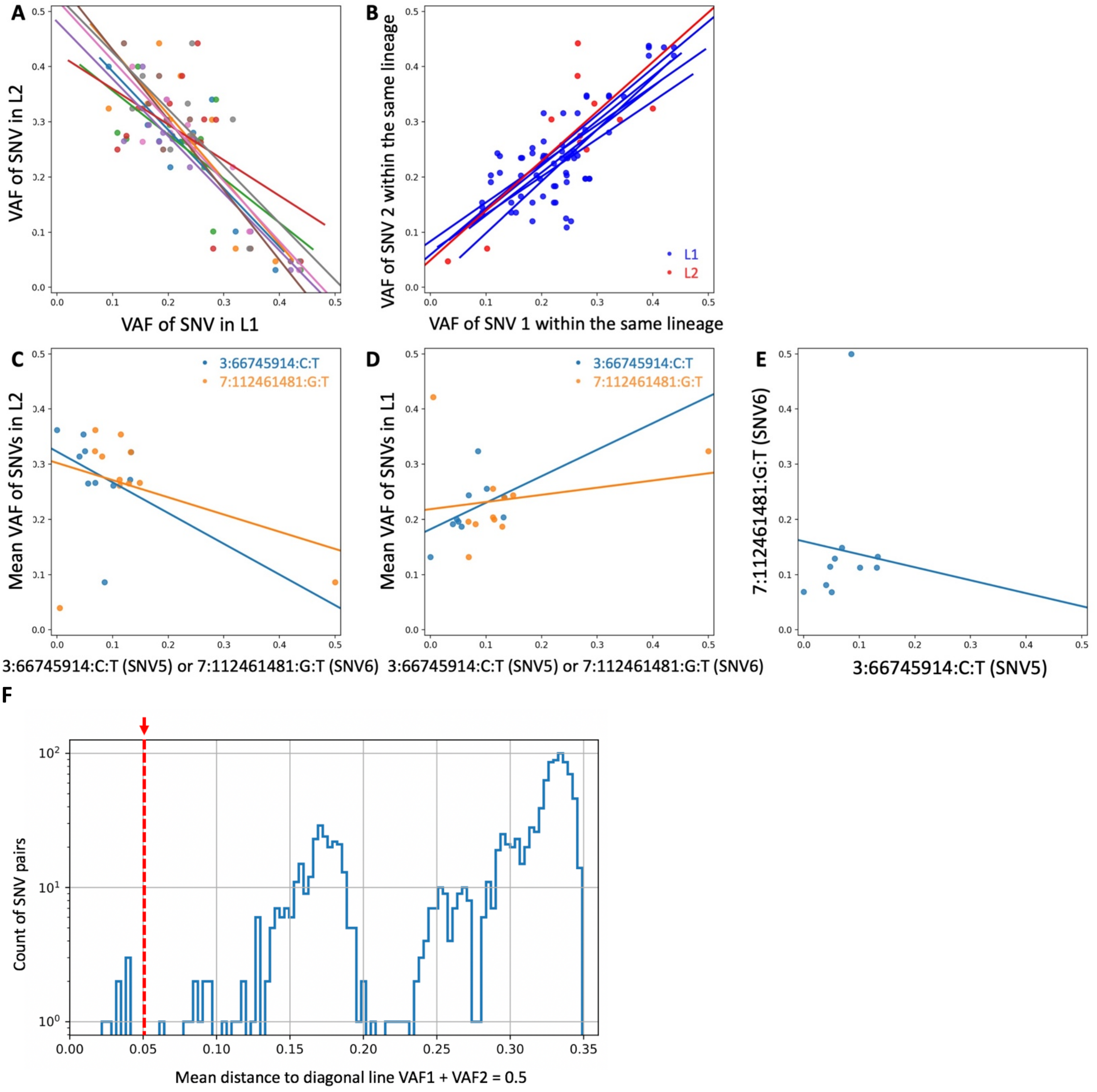


**Supplementary Figure 11: Cell lineage analysis from multiple bulks.** *(A)* Each color corresponds to a pair of mosaic SNVs with one in the L1 (x-axis) and another in L2 (y-axis) lineages (see **Fig. 4**). Each dot represents VAF values for a SNV pair in a sample. The lines indicate VAF linear regression plots across multiple samples for a given pair of SNVs. *(B)* Mosaic SNVs defining the same cell lineage (either L1 or L2) show correlated VAFs across multiple tissue samples. *(C)* The VAFs of mosaic SNVs defining L1 sub-lineages (x-axis, SNV5 and SNV6) are anti-correlated with the VAFs of the SNVs in the L2 lineage (y-axis). *(D)* The VAFs of mosaic SNVs defining L1 sub-lineages (x-axis, SNV5 and SNV6) are positively correlated with the mean VAFs of mosaic SNVs in the L1 lineage (y-axis). *(E)* The VAFs of SNV5 and SNV6 defining different L1 sub-lineages are anti-correlated across multiple samples. (*F)* The distribution of all possible pairs of 49 mosaic SNVs by the distance to a diagonal line in VAF space (aka in Figure 4).

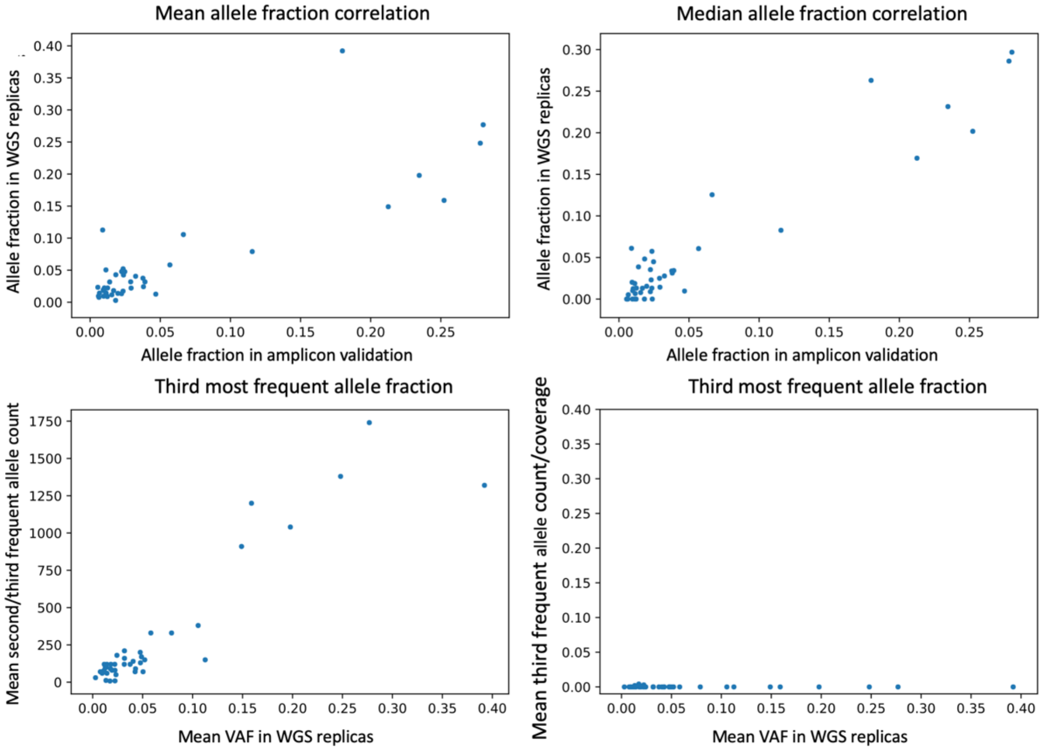


**Supplementary Figure 12: Summary statistics for validated mosaic SNVs.** Each dot represents a mosaic SNV. Top: Mean (left) and median (right) somatic SNV VAFs detected in the PCR amplicon-based sequencing validation experiments (x-axis) positively correlate with their respective VAFs in WGS datasets (y-axis). Bottom left: Plotted are the mean VAF of a mosaic variant in WGS replicates *vs.* the mean allele counts (across replicates) of the second most frequent allele (Rank 2) divided by the mean count of third most frequent allele (Rank 3). Each dot corresponds to the site of a validated mosaic SNV. Rank 2 represents the VAF of the mosaic allele and Rank 3 is a proxy for the background signal. As expected, supporting evidence for somatic SNVs in WGS datasets increase as a function of their VAF. Bottom right: Plotted are the mean VAF of a mosaic variant in WGS replicates *vs.* the mean count of the third most frequent allele divided by the sequence coverage. The resultant values are independent of their VAF, which is consistent with Rank 3 being a proxy for the background signal.

**Supplementary Table 1: Summary of results from the six analytical methods used to call mosaic SNVs.** Columns 1 and 2: The number of candidates identified by each calling method. Columns 3 to 6: Types of SNVs identified using each Method (see Table 1). Column 7: The set of 400 candidate mosaic SNVs selected for validation experiments, with number validated in parenthesis. The last row presents the total non-redundant counts across the methods, collapsing overlapping candidates into single events.

| | Total Candidates | Absolute Singletons (validated/attempted) | Data Source Singletons (validated/attempted) | Approach Singletons (validated/attempted) | Multi-calls (validated/attempted) | Attempted for validation (validated) |
|---|---|---|---|---|---|---|
| Method 1 | 13 | 1 (0/1) | 0 (0/0) | 11 (0/11) | 1 (0/1) | 13 (0) |
| Method 2 | 16 | 0 ((0/0) | 0 (0/0) | 9 (0/9) | 7 (7/7) | 16 (7) |
| Method 3 | 53 | 5 (0/5) | 2 (0/2) | 13 (8/13) | 33 (28/33) | 53 (36) |
| Method 4 | 57 | 3 (0/3) | 1 (0/1) | 24 (0/24) | 29 (19/29) | 57 (19) |
| Method 5 | 100 | 4 (0/4) | 2 (0/2) | 63 (1/63) | 31 (24/31) | 100 (25) |
| Method 6 | 1148 | 135 (0/27) | 3 (0/3) | 979 (1/200) | 31 (22/31) | 258 (23) |
| Total | 1298 | 148 | 4 | 1101 | 45 | 400 (43) |

**Supplementary Table 2**: List of 400 sites for validation (external file)

**Supplementary Table 3: Summary of results from droplet digital PCR validation experiments.** We initially attempted to design ddPCR assays to assess the validity of 29 candidate mosaic SNVs; we successfully designed and conducted validation experiments for 13 candidate mosaic SVNs. Columns 1 and 2: The candidate SNV chromosomal location and its nucleotide position in the GRCh37d5 human genome reference sequence. Columns 3 and 4: The GRCh37d5 (ref) and alternative (alt) alleles. Column 5: Summary of the results from the ddPCR experiments. Column 6: The ratio of ddPCR products containing the alternative allele. Columns 7 and 8: Decision status after completion of the ddPCR experiments.

| ch | pos | ref | alt | ddPCR Result | Ratio | previous decisions | VAF in amplicon seq | final decision |
|---|---|---|---|---|---|---|---|---|
| 3 | 150634574 | G | A | Positive | 2.88% | PASS | 1.28%, 2.06% | PASS |
| 4 | 3857654 | C | T | Positive | 21.85% | PASS | 28.3%, 27.3% | PASS |
| 7 | 112009378 | G | A | Positive | 1.94% | PASS | 1.01%, 2.89% | PASS |
| 14 | 72870518 | A | G | Positive | 19.29% | PASS | 25.23% ,21.67% | PASS |
| 15 | 46636659 | C | G | Positive | 2.67% | PASS | 1.83%, 1.83% | PASS |
| 7 | 80158107 | C | T | Positive | 1.03% | PASS | 0.49%, 1.78% | PASS |
| 11 | 120386729 | C | T | Positive | 0.82% | Validated | 0.010% ,2.47% | PASS |
| 6 | 47565225 | G | C | Positive | 0.86% | Validated | 0.66%, 0.58% | PASS |
| 14 | 35629120 | G | A | Positive | 34.39% | Germline | 32.90%, 33.11% | Germline |
| 5 | 113236393 | C | A | Negative | 0.02% | PASS | 0.53% | false_positive |
| 2 | 81089746 | T | C | Negative | 0.00% | false_positive | 0.02%, 0.003% | false_positive |
| 18 | 12614010 | A | G | Negative | 0.00% | false_positive | 0.98% | false_positive |
| 14 | 97512875 | A | C | Negative | 0.02% | False positive by VAF < 0.5% | 0.12%, 0 | false_positive |

**Supplementary Table 4: The naïve Bayes probability calculation criteria used to evaluate candidate mosaic SNVs.** Column 1: Criteria used to validate somatic SNVs (see below). Column 2: The probability that a given mosaic SNV will be validated (Pr[PASS]) when considering each criterion listed in Column 1. Column 3: The probability that a given mosaic SNV call will not be validated (PR[FAIL]) when considering each criterion listed in Column 1. Larger differences between the probability values listed in Columns 2 and 3 indicate a higher discriminative potential for each criterion.

The criteria used to evaluate candidate somatic SNVs using Chromium 10X linked-read datasets (*10X information*) are: (i) a lack of information using Chromium 10X linked-read data to evaluate a given putative mosaic SNV call (*10X no info*); (ii) a candidate mosaic SNV that is supported using 10X Chromium linked-read data (*10X ok*); and (iii) a candidate mosaic SNV is excluded using the Chromium linked-read data (*10X fail*).

The criteria used to evaluate candidate somatic SNVs by detecting alternative reads in single-cell sequencing datasets (*Single-cell ALT read information*) are: (i) the number of single cells containing between 1 and 6 alternative allele reads for a given candidate mosaic SNV (*0 < # single cells with ALT reads < 6*); (ii) none of the single cells contain alternative allele reads for a given candidate mosaic SNV (*No single cells have ALT reads*); and (iii) the number of single cells containing >6 alternative reads for a given candidate mosaic SNV *(> 6 single cells have ALT reads*).

The criteria used to evaluate candidate somatic SNVs using alternative reads using the single-cell genotyper (***Single cell genotype information*) are:** (i) the single-cell genotyper did not provide sufficient information for the candidate mosaic SNV (*Single cell genotypes inconclusive*); (ii) the single-cell genotyper concluded the candidate mosaic SNV was poorly amplified in all single cells (*Position is poorly amplified*); and (iii) the single-cell genotyper yielded evidence supporting a candidate mosaic SNV call in one or more cells (*Evidence of SNV in single cells*).

The categories used to evaluate candidate somatic SNVs using the 1000 Genomes Project PON filter (***PON information***) are: (i) five or fewer individual genomes in the 1000 Genomes Project have alternative allele reads corresponding to the candidate somatic SNV call (*5 or fewer PON samples have ALT reads*); and (ii) more than five individual genomes in the 1000 Genomes Project have alternative allele reads corresponding to the candidate somatic SNV call (*>5 PON samples have ALT reads*).

| | **Pr(PASS)** | **Pr(FAIL)** |
|---|---|---|
| **10X information** | | |
| 10X no info | 0.435 | 0.544 |
| 10X ok | 0.500 | 0.286 |
| 10x fail | 0.065 | 0.169 |
| **Single cell ALT read information** | | |
| 0 < # single cells with ALT reads < 6 | 0.391 | 0.525 |
| No single cells have ALT reads | 0.565 | 0.336 |
| > 6 single cells have ALT reads | 0.043 | 0.139 |
| **Single cell genotype information** | | |
| Single cell genotypes inconclusive | 0.711 | 0.904 |
| Position is poorly amplified | 0.022 | 0.006 |
| Evidence of SNV in single cells | 0.267 | 0.089 |
| **PON information** | | |
| 5 or fewer PON samples have ALT reads | 0.800 | 0.136 |
| >5 PON samples have ALT reads | 0.200 | 0.864 |

**Supplementary Table 5**: List of amplicon sequencing primers for the 400 sites of validation (external file).

**Supplementary Table 6**: List of primers and probes for sites performed ddPCR (external file).
